# The action of physiological and synthetic steroids on the calcium channel CatSper in human sperm

**DOI:** 10.1101/2023.05.10.540165

**Authors:** Lydia Wehrli, Ioannis Galdadas, Lionel Voirol, Martin Smieško, Yves Cambet, Vincent Jaquet, Stéphane Guerrier, Francesco Luigi Gervasio, Serge Nef, Rita Rahban

## Abstract

The sperm-specific channel CatSper (**cat**ion channel of **sper**m) controls the intracellular Ca^2+^ concentration ([Ca^2+^]_i_) and plays an essential role in sperm function. It is mainly activated by the steroid progesterone (P4) but is also promiscuously activated by a wide range of synthetic and physiological compounds. These compounds include diverse steroids whose action on the channel is so far still controversial. To investigate the effect of these compounds on CatSper and sperm function, we developed a high-throughput-screening (HTS) assay to measure changes in [Ca^2+^]_i_ in human sperm and screened 1,280 approved and off-patent drugs including 90 steroids from the Prestwick chemical library. More than half of the steroids tested (53%) induced an increase in [Ca^2+^]_i_ and reduced the P4-induced Ca^2+^ influx in human sperm in a dose-dependent manner. Ten of the most potent steroids (activating and inhibiting) were selected for a detailed analysis of their action on CatSper and their ability to act on sperm motility, acrosomal exocytosis (AR), and penetration in viscous media. We found that these steroids show an inhibitory effect on P4 but not on prostaglandin E1-induced CatSper activation, suggesting that they compete for the same binding site as P4. Pregnenolone, dydrogesterone, epiandrosterone, nandrolone, and dehydroepiandrosterone acetate (DHEA) were found to activate CatSper at physiological concentrations. Stanozolol, epiandrosterone, and pregnenolone induced AR similarly to P4, whereas stanozolol and estropipate induced an increase in sperm penetration into viscous medium. Furthermore, using a hybrid approach integrating pharmacophore analysis and statistical modelling, we were able to screen *in silico* for steroids that can activate the channel and define the physicochemical and structural properties required for a steroid to exhibit agonist activity against CatSper. Overall, our results indicate that not only physiological but also synthetic steroids can modulate the activity of CatSper with varying potency and affect human sperm functions *in vitro*.

## Introduction

In order to fertilize an oocyte, sperm must first undergo multiple processes such as capacitation, acquisition of hyperactivation, and acrosomal exocytosis, which are known to be controlled in part by an increase in intracellular calcium concentration ([Ca^2+^]_i_). The influx of calcium ions in human sperm is primarily mediated by the sperm-specific cationic channel of sperm (CatSper) (Ren and Xia, 2010). CatSper is activated by alkalization of intracellular pH (pH_i_), membrane depolarization (Kirichok et al., 2006), and by natural ligands such as the steroid progesterone (P4) and the arachidonic acid derivative prostaglandins, mainly prostaglandin E1 (PGE1) (Lishko et al., 2011, Strünker et al., 2011). These ligands have distinct binding sites to activate the channel (Strünker et al., 2011) and it has been suggested that P4 does not directly activate CatSper but rather binds to an upstream lipid hydrolase called the α/β hydrolase domain-containing protein 2 (ABHD2) (Miller et al., 2016). Upon P4 binding, ABHD2 degrades the endocannabinoid 2-arachidonoylglycerol (2-AG), which blocks the channel and thereby releases CatSper from inhibition (Miller et al., 2016). Studies over the last 30 years have shown that various endogenous and exogenous steroids are also able to evoke Ca^2+^ influx and motility responses in human sperm (Blackmore et al., 1990, Blackmore et al., 1996, Rossato et al., 2005, Luconi et al., 2004, Luconi et al., 1999, Brenker et al., 2018a, Brenker et al., 2018b, Rehfeld, 2020, Jeschke et al., 2021). Steroids such as 17-OH-progesterone, estradiol, dihydrotestosterone, androstenedione, and pregnenolone have been described to act via CatSper (Brenker et al., 2018a, Brenker et al., 2018b, Jeschke et al., 2021, Lishko et al., 2011, Rehfeld, 2020, Strünker et al., 2011). However, some of the results are still controversial. For example, testosterone, estradiol, and hydrocortisone have been reported to be both CatSper antagonists (Mannowetz et al., 2017) and agonists (Brenker et al., 2018b, Rehfeld, 2020). Furthermore, several studies demonstrated that CatSper is promiscuous and can be activated by a wide range of synthetic or natural compounds such as odorants (Spehr et al., 2003), endocannabinoids (Miller et al., 2016), and Endocrine Disrupting Chemicals (EDCs) (Schiffer et al., 2014, Rehfeld et al., 2016, Tavares et al., 2013). The list of compounds that can activate the channel was recently expanded to include commonly prescribed pharmaceutical drugs such as the selective serotonin reuptake inhibitors (SSRI) class of antidepressants (Rahban et al., 2021), the 5α reductase inhibitor finasteride (Birch et al., 2021), as well as paracetamol metabolite N-arachidonoyl phenolamine (AM404) (Rehfeld et al., 2022). These results are consistent with the finding that human CatSper serves as a polymodal chemosensor that harbors promiscuous binding sites for structurally diverse ligands (Brenker et al., 2012). However, the physiological effects on sperm cells and the molecular characteristics that allow multiple structurally diverse steroids, whether physiological or synthetic, to activate CatSper remain poorly understood and contradictory.

Using our novel high-throughput screening (HTS) assay, we have developed an accurate and rapid method to investigate the effects of a wide range of pharmaceutical drugs, including synthetic and physiological steroids, on CatSper. We studied in detail ten steroids that can directly activate the channel and interfere with the P4 but not the PGE1-induced calcium response. We also evaluated the effect of these ten steroids on the ligand-independent activation of the channel through intracellular alkalization and their effects on acrosomal exocytosis and penetration into viscous media. Finally, pharmacophore and statistical modelling were used to screen *in silico* for steroids that can activate the channel and better understand the molecular characteristics of the steroids that promote CatSper activation.

## Materials and Methods

### Reagents

The PRESTWICK library composed of 1,280 compounds was purchased by the R.E.A.D.S. unit of the University of Geneva. The drugs were dissolved at a stock concentration of 10 mM in dimethyl sulfoxide (DMSO). Progesterone and PGE1 were purchased from Sigma-Aldrich (Buchs, Switzerland). The fluorescent Ca^2+^ indicator Fluo-4 AM was purchased from Invitrogen and dissolved in DMSO at a final concentration of 2.5 µM (California, USA). The nuclear dye Propidium iodide (PI) was purchased from Thermo Fisher Scientific (MA, USA), diluted in water at stock concentration of 1mg/mL. Pisum sativum agglutinin (PSA) staining dye, was purchased from from Sigma-Aldrich (Buchs, Switzerland). Human serum albumin (HSA) was obtained from Polygon Diagnostics (Lucerne, Switzerland). MDL 12330A hydrochloride was purchased from Tocris Bioscience (Bristol, UK). The following steroids were purchased from Sigma-Aldrich (Buchs, Switzerland): progesterone, dehydroisoandosterone 3-acetate (DHEA), estropipate, stanozolol, epiandrosterone, deoxycorticosterone, dydrogesterone, 3-alpha-Hydroxy-5-beta-androstan-17-one, pregnenolone, medrysone, equilin, estrone, norgestimate, canrenoic acid potassium salt, exemestane, finasteride, lithocholic acid, formestane, exemestane, fluorometholone, estradiol-17 beta, androsterone, medrysone, nandrolone, fluorometholone, oxymetholone, urosiol, oxandrolone, canrenone, fulvestrant, lynestrenol, norethindrone, ethisterone, nomegestrol acetate, gestrinone, chlormadinone acetate, norgestrel-(-)-D, ethynodiol diacetate, mifepristone, budesonide, adrenosterone, ethinylestradiol, megestrol acetate, ethynylestradiol 3-methyl ether, 3α,21-dihydroxy-5α-pregnan-20-one, 5α-pregnan-3α-ol-20-one, 19-norandrosterone, androsterone, 5alpha-pregnan-20-one, 5alpha-androstan-3-one, 5alpha-pregnane-3,20-dione, androstenedione, tanaproget, BB_NC-00182 and dissolved at a stock solution of 20 mM in DMSO. To implement the pharmacophore model, the steroids were purchased from ChemDIV (California, USA) under the following ID numbers: N017-0003, N039-0025, N050-0010, 3647-1627, N044-0005, 0449-0076, 2361-0084, N039-0043, N050-0013, N039-0021, 0449-0077, N017-0007, 8011-0433, N050-0008, 3529-0011, 3529-0046, 5263-0009, N050-0018, and 3β,21-dihydroxy-5α-pregnan-20-one, 21-hydroxypregnenolone and MolPort-001-728-767 were purchased from MolPort (Riga, Latvia), and promegestrone was purchased from BOC Sciences (NY, USA) and dissolved at a 20 mM stock concentration in DMSO.

### Sample collection and preparation

Human semen samples were collected from healthy donors by masturbation after a recommended sexual abstinence period of minimum of two days. Semen samples were allowed to liquefy at 37°C for 15-30 minutes. Motile spermatozoa were selected by a ‘swim-up’ procedure as previously described (Strünker et al., 2011). Briefly, 1 mL of semen was deposited in a 45°-inclined tube with 4 mL of Human Tubular Fluid (HTF) medium containing: 93.8 mM NaCl, 4.7 mM KCl, 0.2 mM MgSO_4_, 0.37 mM KH_2_PO_4_, 2.04 mM CaCl_2_, 20.98 mM HEPES, 2.78 mM glucose, 21.4 mM Na-lactate, 4 mM NaHCO_3_, 0.33 Na-pyruvate, and the pH was adjusted to 7.35. Motile sperm were allowed to swim at 37°C for one hour. The supernatant was collected in 15 mL Falcon tubes and washed twice by centrifugation at 700 x g for 20 minutes. Sperm were supplemented with 3 mg/mL HSA. Written informed consent was obtained from all donors prior to donation and approved by the Cantonal Ethics Committee of the State of Geneva (CCER #14-147).

### High throughput measurements of changes in [Ca^2+^]_i_

Cells were loaded with the fluorescent Ca^2+^ indicator Fluo4-AM at a final concentration of 2.5 µM for 30 minutes at 37°C. Excess dye was washed via centrifugation at 700 x g for 20 minutes and sperm were resuspended in HTF medium to obtain a final concentration of 6.25 × 10^6^ cells/mL. 20 μL of cells loaded with Fluo4-AM were dispensed in a 384-wells plate (Corning, NY, USA) to allow fluorescence measurement. The change in [Ca^2+^]_i_ was assessed simultaneously in all wells using a Functional Drug Screening System (FDSS/μCELL) (Hamamatsu Phototonics K.K., Yokohama, Japan) at 37°C. The dye was excited at 530 nm, and emission was recorded at 480 nm. Fluorescence was measured before and after the addition of 5 µL of compounds, vehicle control (DMSO 0.5%), positive control (P4 or PGE1 at 2 µM or NH_4_Cl at 10 mM for alkalization-induced calcium measurement), and negative control (MDL at 20 µM). Fluorescence was recorded for a total of 16.5 minutes: 30 seconds before injection, 12 minutes after injection, and 4 minutes after the second injection. The Prestwick library was tested in technical duplicate. Calcium response curves were normalized using the following formula, 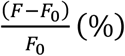, where *F*_*0*_ is the average of the basal fluorescence prior to the addition of the compound of interest and *F* corresponds to the fluorescence value at each time point. The ΔF/F_0_ of the vehicle control (DMSO 0.5%) was subtracted from all the other values to adjust for dilution and pipetting artifacts. All compounds were normalized to their respective controls. Positive hits were selected when all four parameters showed a coefficient of variance (CV) of less than 20%. To generate sigmoidal curves, the concentrations were converted to their respective log values, and the data were normalized using the following formula: 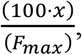, where F_max_ is the maximum value of the positive control. A four-parameter Hill equation with a variable Hill coefficient was fitted to generate the EC_02,_ EC_50,_ and IC_50_. Data were plotted and analysed using GraphPad Prism, version 8.1.1.

### Assessment of acrosomal exocytosis

The evaluation of acrosomal exocytosis in live human sperm cells was performed using a modified version of a previously described method (Zoppino et al., 2012). Briefly, sperm cells were allowed to capacitate for 4 hours at 37°C in capacitating media (HTF+) containing: 93.8 mM NaCl, 4.7 mM KCl, 0.2 mM MgSO_4_, 0.369 mM KH_2_PO_4_, 2.04 mM CaCl_2_, 20.98 mM HEPES, 2.78 mM glucose, 21.4 mM Na-lactate, 25 mM NaHCO_3_, 6.6 µM pyruvate, and the pH was adjusted to 7.35 with NaOH. Capacitated sperm were then incubated for 30 minutes with 5 µg/mL *Pisum sativum* agglutinin conjugated with fluorescein isothiocyanate (PSA-FITC) and 10 µg/mL Hoechst-33342. Samples were centrifuged at 700 x g for 10 minutes at room temperature and resuspended in HTF+ supplemented with 0.5 µg/mL PI. Sperm were then incubated for 45 minutes in the absence (DMSO) or presence of the steroids at 10 µM, along with positive controls P4 at 10 µM. The ionophore A23187 was incubated at 2 µM for 30 minutes and samples were analyzed using BDFACSAria. PSA-FITC positive and PI negative cells were classified as live acrosome-reacted cells, whereas PSA-FITC negative and PI negative cells were classified as live acrosome intact cells. Doublet exclusion was performed by analyzing two-dimensional dot plot forward-scatter width (FSC-W) against forward-scatter height (FSC-H), and by analyzing side-scatter width (SSC-W) against side-scatter height (SSC-H) on the pre-selected singlets. Data were collected from 40,000 PSA-FITC negative events per sample to define the sperm population. Based on the selected population, PSA-FITC-positive cells were selected as acrosomal reacted cells, whereas PSA-FITC-negative cells were selected as acrosome intact cells.

### Sperm motility assessment

Sperm motility was assessed using a computer-assisted sperm analysis (CASA, Hamilton Throne Ceros II, UK). Swim-up prepared sperm cells were diluted to a concentration of 12.5 × 10^6^/mL in HTF+ and were allowed to capacitate for 3 hours at 37°C in HTF+. Capacitated sperm were then incubated for an additional hour in the absence (DMSO) or presence of steroids of interest, progesterone, or PGE1 at 10µM. A 2 µL aliquot was placed in a counting chamber with a 20 µm depth (Leja, Nieuw-Vennep, The Netherlands) and a minimum of 200 spermatozoa in at least five randomly selected fields were captured. Progressive motility was classified based on the average path velocity (VAP) with the following parameters: slow <1 > medium < 25 > rapid. Hyperactive cells were classified based on the following parameters: VAP > 50 µm/s, a curvilinear velocity (VCL) > 90 µm/s, and an amplitude of lateral head (ALH) > 5 µm.

### Sperm penetration in viscous media

The ability of sperm to penetrate in viscous media similar in viscosity to that found in the female reproductive tract was assessed using the modified Kremer test, as previously described (Rahban et al., 2021). Briefly, sperm cells at 3 × 10^6^/mL were incubated in the absence (DMSO) or presence of steroids at 10 µM for 1 hour at 37°C. A glass capillary (0.2 × 4.0 × 50 mm, CM scientific, UK) was filled with 1% (w/v) methylcellulose (MC, 4000 centipoises) prepared with HTF media supplemented with 3mg/mL HSA and was sealed on one end with wax (Vitrex, UK). The glass capillary tube was then added to the sperm cells on the open end. Progesterone (10 µM), steroids (10 µM), or DMSO was also added to the MC in the glass capillary tubes. Sperm penetration was assessed after 1 hour of incubation at 37°C by counting sperm at 1 cm using the CASA with a 10x objective.

### Pharmacophore and statistical modelling

In order to build a ligand-based model that could be used to screen *in silico* for additional steroid-based compounds with an agonist effect, we started by constructing an initial pharmacophore model and a semi-parametric zero-inflated beta regression model using the 90 steroids identified through the screening of the Prestwick library (see **Supplementary data** for the details on the construction of the models). Given the known importance of different physicochemical properties of compounds for their binding to charged porins in bacteria, including their polarity (Acosta-Gutierrez et al., 2018), we developed a physicochemical property-based filtering criterion relying on a statistical regression model. This model was then used in combination with our pharmacophore model to increase the probability of the *in silico* screening. The regression model assumes a zero-inflated beta distribution to model the agonist effect, which is the dependent variable, while it considers the original physicochemical properties and their second and third-order interactions between them as independent variables. The QikProp tool of the Schrödinger suite (QikProp, Schrödinger, LLC, New York, NY, 2021) was used to calculate the physicochemical properties of each compound. Of the 10 physicochemical properties selected using the Adaptive Lasso procedure (Zou, 2006) and included in the model, seven were identified as significant when considering a confidence level of 1 − α = 0.95, most of which were the result of interaction between variables. Out of these variables, only one was an original variable, while the rest of the significant variables were interactions between two or three initially measured properties. To select the most optimal pharmacophore and statistical model combination, we used a blind dataset of 18 compounds whose agonist effect was predicted by the pharmacophore and statistical model prior to their experimental testing. Among the different pharmacophore models considered, the one with the two hydrophobic sites and one acceptor site, combined with a beta zero-inflated model with adaptive-Lasso selected covariates was considered the most optimal (**Supplementary Fig. 1**). In particular, the combination of the two models was able to identify five true positives and one false positive out of the six compounds. The results of the blind test demonstrated that the combination of the two models was crucial to achieving a good prediction ratio, as considering either model alone led to several false positives. Using this combination of models, we screened *in silico* the GeDB (n=187), ChemDiv (n=498), and ZINC (n=98) libraries after querying only for compounds with a steroidal core. For all the compounds screened virtually, 30 additional steroids were tested: the top 30 with the highest pharmacophore fitting score and predicted agonist effect, and five that were expected to have no effect but were included for further validation of the model were purchased and tested experimentally.

## Results

### The Prestwick library screen and confirmation of hits by dose-response

To evaluate the potential effect of synthetic and physiological compounds including steroids on calcium influx, we screened the Prestwick library which consists of 1,280 compounds from various therapeutic classes. Our HTS assay was designed to screen for compounds that can induce a calcium influx (screen 1, activators) and/or can alter the P4- and PGE1-induced calcium influx (screen 2, inhibitors) (**Fig. 1, A**). Compound plates were validated with a Z’-factor > 0.4 (**Fig. 1, B**). Hit selection was based on an arbitrary cutoff of 30 % activation and/or inhibition compared to the controls (**Fig. 1, C**). We identified 106 (8.3%) hits, of which 31 were putative activators, 45 were putative inhibitors, and 30 had a dual effect (**Fig. 2, A**). Interestingly, the compounds with dual activating and inhibiting effects were mostly steroids. All 26 of them reduced the P4-but not PGE1-induced calcium influx (**Fig. 2, B & C**). The hits steroidal compounds were validated by dose-response curves (DRC) in technical duplicates and biological triplicates and their EC_50_s as well as their IC_50_s for the P4-induced response were measured (**Supplementary Table 1**). Interestingly, of the 90 steroids in the Prestwick library, more than half (53%) were able to both activate and inhibit P4-induced calcium influx in human sperm (**Fig. 2, B**). In contrast, inhibition of PGE1-induced calcium influx was rarely observed with these steroids (**Fig. 2, C**). For the next part of this study, the 10 steroids with the strongest dual activation and inhibition potential were selected for further investigation. These were the synthetic steroids stanozolol, estropipate, dydrogesterone, medrysone, and the natural steroids dehydroepiandrosterone acetate (DHEA), testosterone, epiandrosterone, nandrolone, pregnenolone, deoxycorticosterone.

**Figure 1:**
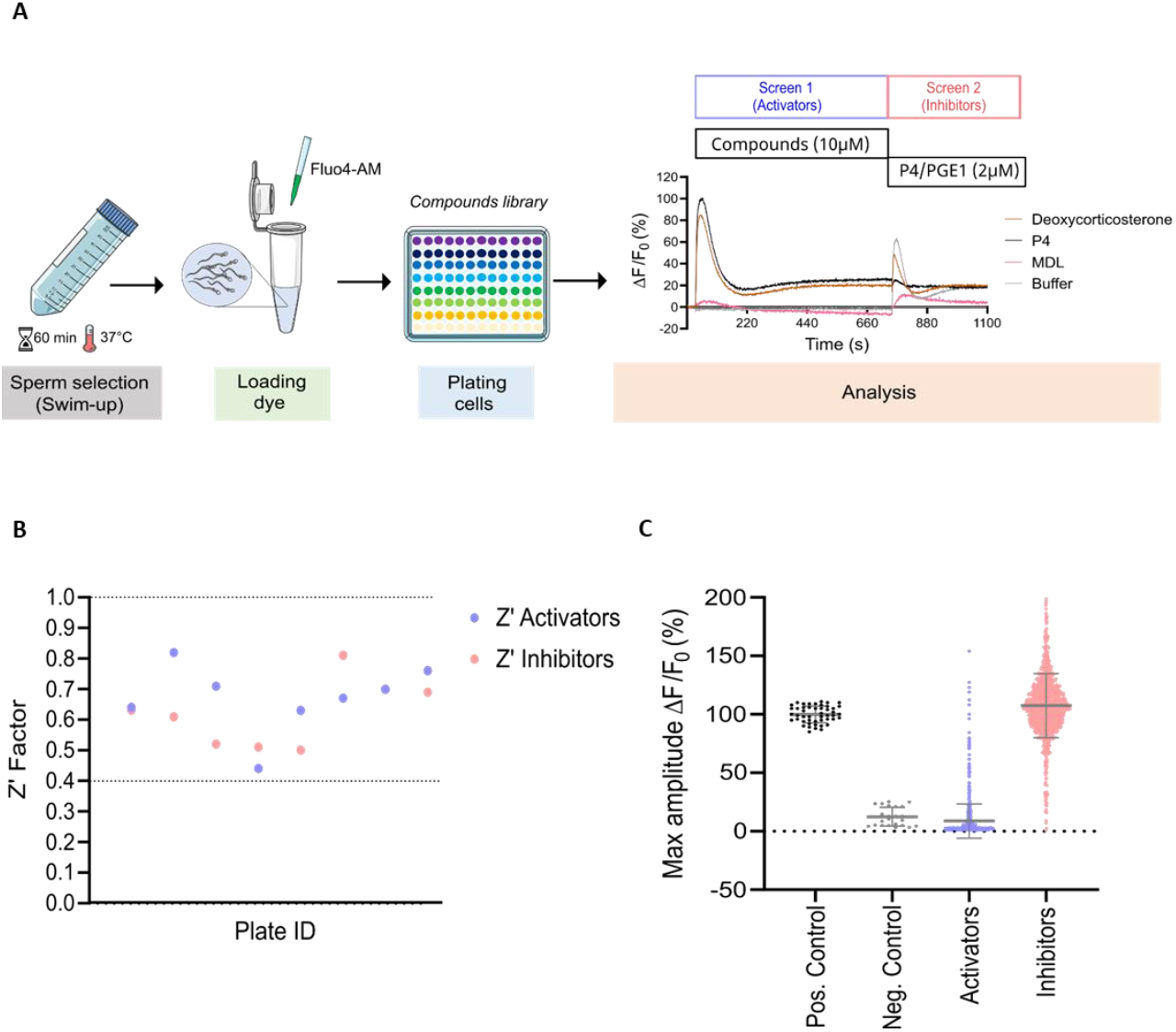
Prestwick library screen in human spermatozoa. (**A**) Graphical overview of the workflow used to screen the Prestwick library composed of 1280 compounds that was designed to screen for molecules capable of inducing calcium influx (screen 1, activators) and/or altering progesterone (P4)- or prostaglandin E1 (PGE1) -induced calcium influx (screen 2, inhibitors). Representative traces of Ca^2+^ influx by deoxycorticosterone (10 µM), P4 (2 µM), non-specific CatSper inhibitor MDL (at 20 µM) used as a negative control and DMSO was used as a vehicle control (buffer). (**B**) The standard high-throughput screening (HTS) metric, Z’ Factor, was used as an indicative measure of assay robustness and to determine assay performance for all screening plates. Dashed lines indicate min/max Z’ values. Screening results are shown in (**C**): each point represents the maximum amplitude of individual compounds tested at 10 µM. Grey lines represent mean ± standard deviations.

**Figure 2:**
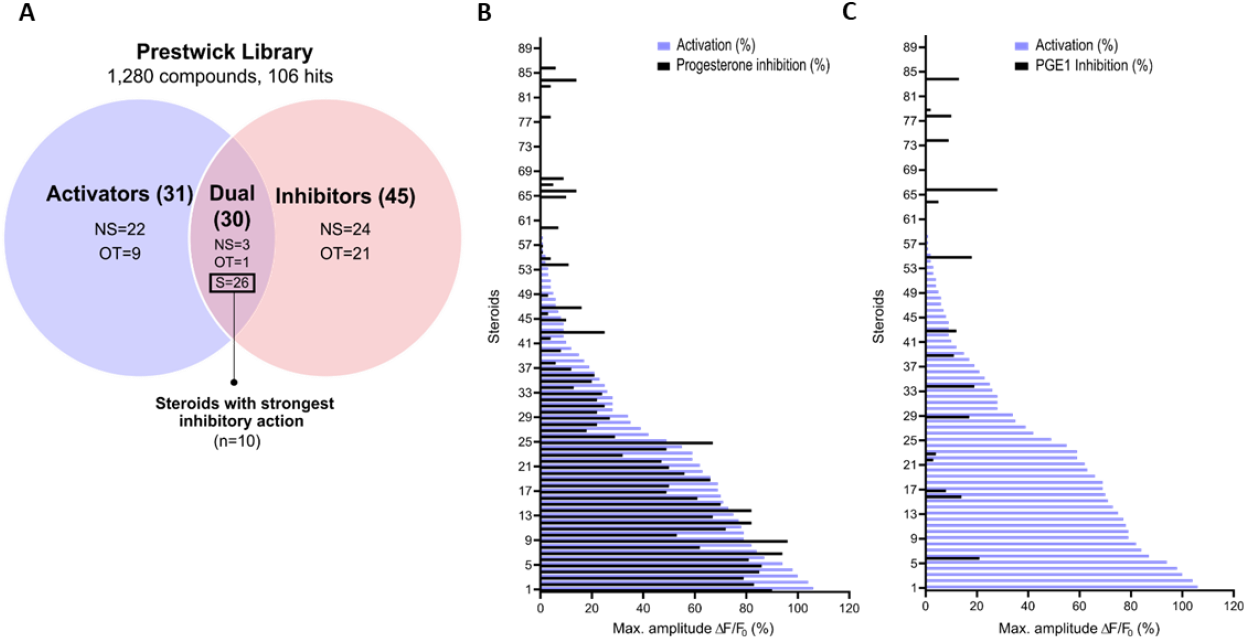
Steroid screening results and steroid selection workflow. (**A**) Schematic representation of the HTS workflow and the selection of steroids. NS: non-steroids; S: steroids; OT: off-targets. Graphical representation of the 90 steroids for their ability to activate or inhibit in (**B**) the P4- and in (**C**) the PGE1-induced Ca^2+^ signaling response. Data are expressed as the mean percentage of maximal amplitude ΔF/F0, triggered by each steroid at 10 µM and normalized to P4 response at 2 µM, before (activation, blue) or after the addition of P4 or PGE1 (inhibition in black).

### Structurally diverse steroids modulate human CatSper

The ten selected steroids were classified into four classes: androgens, estrogens, progestagens, and others, and they all induced an increase in [Ca^2+^]_i_ similar to P4 with estropipate inducing the highest increase in [Ca^2+^]_i_ (**Fig. 3, A & D**). All ten steroids were tested in dose-response experiments and revealed a dose-dependent increase in [Ca^2+^]_I_ (**Supplementary Fig. 2**). More specifically, estropipate, pregnenolone, and dydrogesterone induced a dose-dependent increase in [Ca^2+^]_i_ with values at concentrations as low as 10 nM. Stanozolol and nandrolone also induced a 30% increase in [Ca^2+^]_i_ at a low concentration within the nM range (**Fig. 3 & Supplementary Fig. 2**). EC_50_ values ranged between 0.073 µM for pregnenolone and 0.5 µM for DHEA (**Table 1**). To generate the lowest activating concentration, the EC_02_ was measured for all steroids, and compared with the reported maximum concentration found in blood serum (C_max_), as well as the reported average concentration of steroids found in the follicular fluid (FF) (**Fig. 4**). This comparison highlighted that the steroids progesterone, pregnenolone, dydrogesterone, epiandrosterone, nandrolone, and DHEA act on CatSper within pharmacological concentrations. In the case of nandrolone, progesterone, and dydrogesterone, the EC_02_ values were more than 100 times lower than the C_max_ reported value in the literature (**Table 1** and **Fig. 4**). All steroids tested significantly reduced the P4 response with IC_50_ values ranging between 0.19 µM and 1.75 µM (**Supplementary Fig. 2 & Table 1**). This indicates that P4 is more potent than any of the steroids tested. Interestingly, a strong correlation is observed between the steroids inducing the highest increase in [Ca^2+^]_i_ and the ones that reduce the P4 response the most potently (**Supplementary Fig. 3**). This reduction was not class-specific suggesting that different functional groups can be accommodated in the binding site. Steroids did not alter the PGE1-induced increase in [Ca^2+^]_i_ suggesting that steroids act on the same binding site as that of progesterone but not that of PGE1 (**Fig. 3C** and **F**).

**Table 1:**
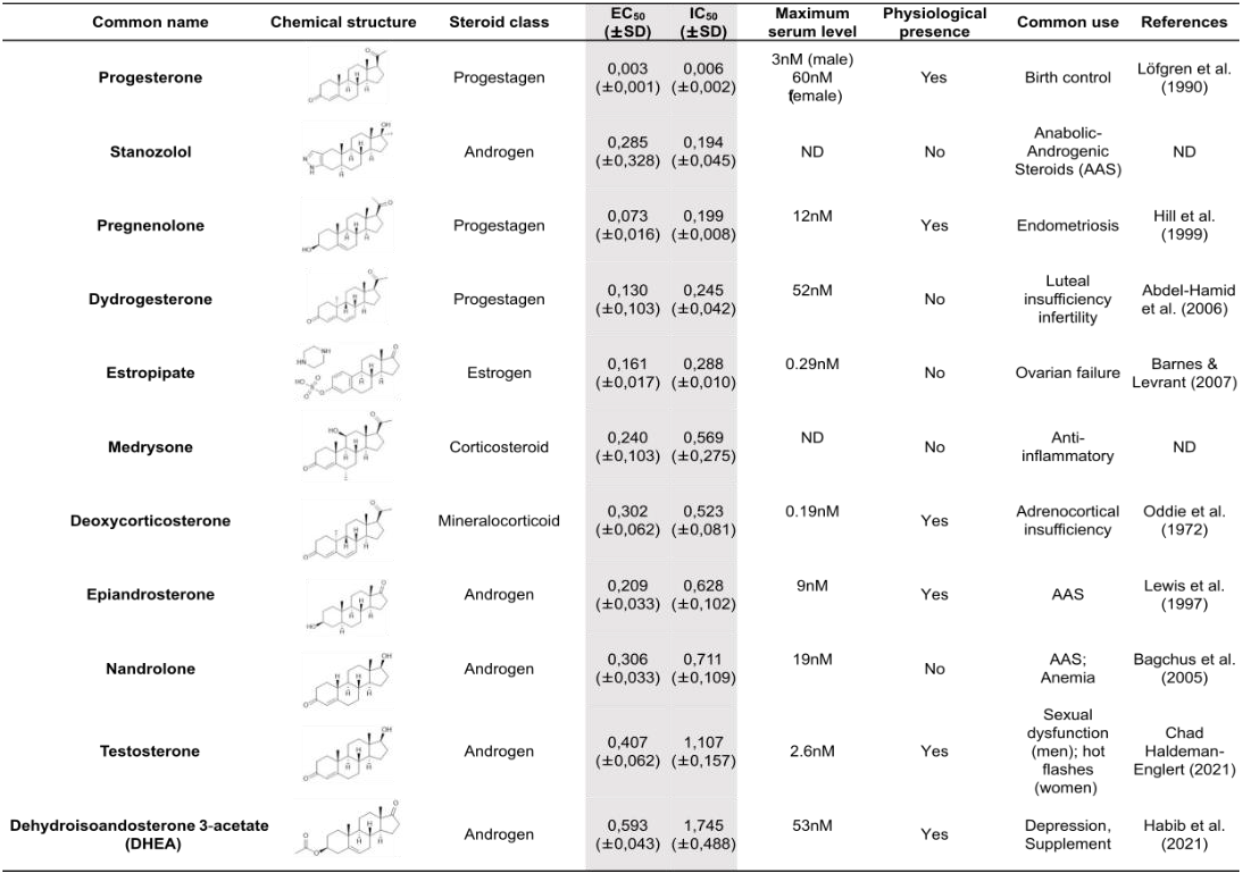
Summary table of EC50 and IC50 values of the ten selected steroids. All EC50 and IC50 were generated following dose-response experiments in absence (DMSO) or presence of steroids ranging from 90 µM to 0.3 nM. EC50/IC50 units are in µM. ND: No data available.

| Common name | Chemical structure | Steroid class | EC<sub>50</sub><br>(±SD) | IC<sub>50</sub><br>(±SD) | Maximum<br>serum level | Physiological<br>presence | Common use | References |
| --- | --- | --- | --- | --- | --- | --- | --- | --- |
| **Progesterone** | | Progestagen | 0,003<br>(±0,001) | 0,006<br>(±0,002) | 3nM (male)<br>60nM (female) | Yes | Birth control | Löfgren et al.<br>(1990) |
| **Stanozolol** | | Androgen | 0,285<br>(±0,328) | 0,194<br>(±0,045) | ND | No | Anabolic-Androgenic<br>Steroids (AAS) | ND |
| **Pregnenolone** | | Progestagen | 0,073<br>(±0,016) | 0,199<br>(±0,008) | 12nM | Yes | Endometriosis | Hill et al.<br>(1999) |
| **Dydrogesterone** | | Progestagen | 0,130<br>(±0,103) | 0,245<br>(±0,042) | 52nM | No | Luteal<br>insufficiency<br>infertility | Abdel-Hamid<br>et al. (2006) |
| **Estropipate** | | Estrogen | 0,161<br>(±0,017) | 0,288<br>(±0,010) | 0.29nM | No | Ovarian failure | Barnes &<br>Levrant (2007) |
| **Medrysone** | | Corticosteroid | 0,240<br>(±0,103) | 0,569<br>(±0,275) | ND | No | Anti-<br>inflammatory | ND |
| **Deoxycorticosterone** | | Mineralocorticoid | 0,302<br>(±0,062) | 0,523<br>(±0,081) | 0.19nM | Yes | Adrenocortical<br>insufficiency | Oddie et al.<br>(1972) |
| **Epiandrosterone** | | Androgen | 0,209<br>(±0,033) | 0,628<br>(±0,102) | 9nM | Yes | AAS | Lewis et al.<br>(1997) |
| **Nandrolone** | | Androgen | 0,306<br>(±0,033) | 0,711<br>(±0,109) | 19nM | No | AAS;<br>Anemia | Bagchus et al.<br>(2005) |
| **Testosterone** | | Androgen | 0,407<br>(±0,062) | 1,107<br>(±0,157) | 2.6nM | Yes | Sexual<br>dysfunction<br>(men); hot<br>flashes<br>(women) | Chad<br>Haldeman-Englert (2021) |
| **Dehydroisoandrosterone 3-acetate**<br>**(DHEA)** | | Androgen | 0,593<br>(±0,043) | 1,745<br>(±0,488) | 53nM | Yes | Depression,<br>Supplement | Habib et al.<br>(2021) |

**Figure 3:**
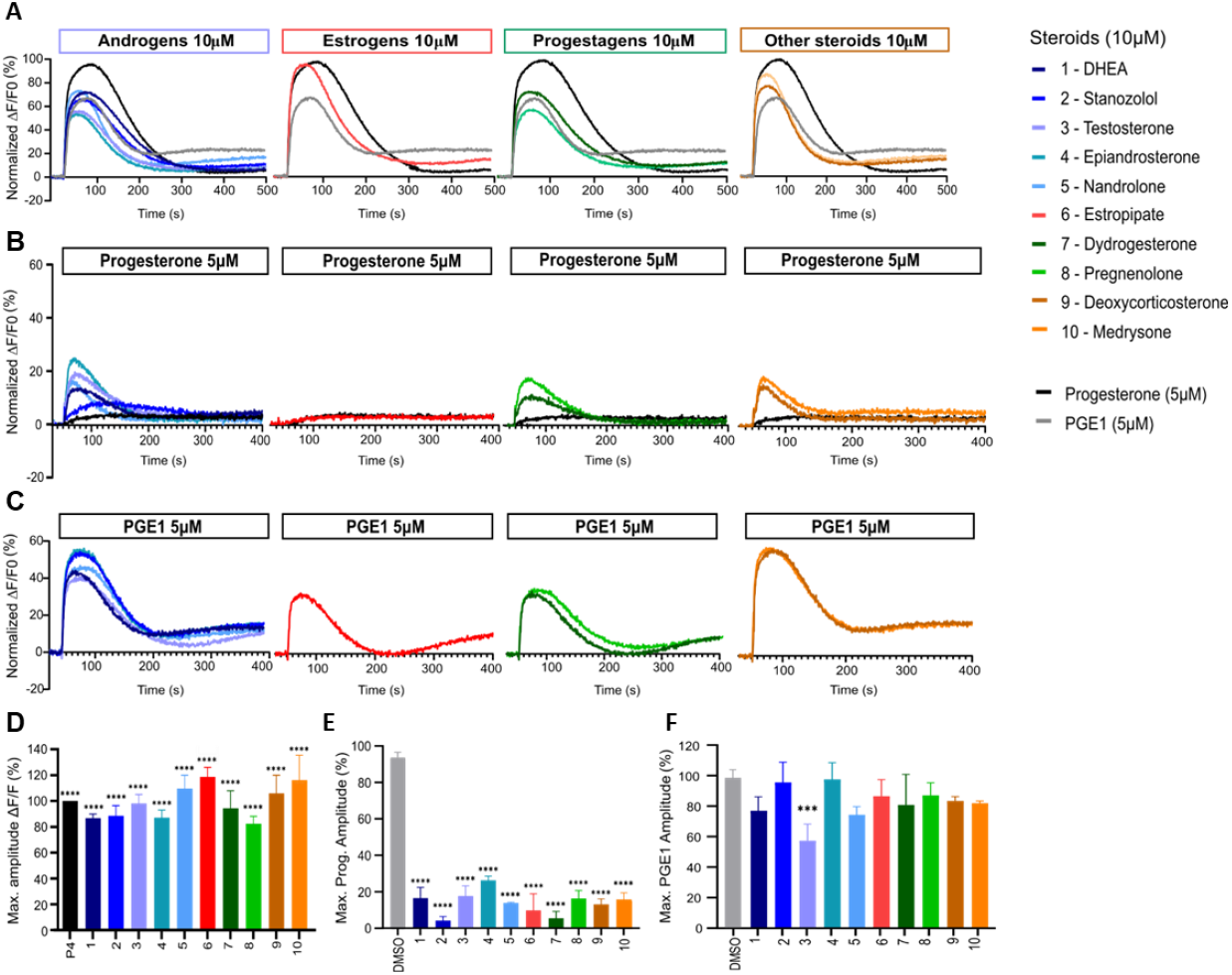
Representative curves of [Ca^2+^]i in human sperm cells induced by diverse steroids. (**A**) Representative Ca^2+^ signals in human sperm evoked by each of the 10 steroids at 10 µM (P4 and PGE1 at 5 µM were used as corresponding positive controls). Ca^2+^ signals evoked in (**B**) by P4 (5 µM) and in (**C**) by PGE1 (5 µM) in the presence of each of the 10 steroids. (**D**) Bar plots representing the maximum amplitude of the increase in [Ca^2+^]i after incubation of the 10 steroids at 10 µM and P4 at 5 µM (n=3). Bar plot showing the maximum amplitude evoked by a single dose of (**E**) P4 (5 µM) and (**F**) PGE1 (5 µM) after incubation with each of the steroids (n=3). p(***)<0.001; p(****)<0.0001.

**Figure 4:**
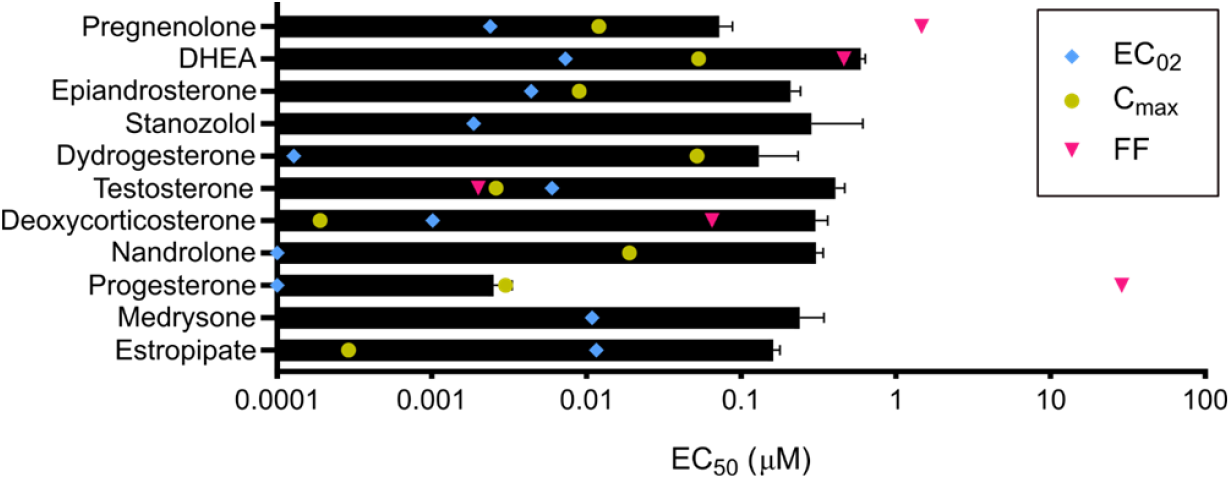
Steroids and their pharmacological concentrations. Half maximal effective concentration (EC50, black) compared with reported maximum serum concentration (Cmax, yellow), average concentration in follicular fluid (FF, pink), and minimum effective concentration (EC02, blue), (n=3). Cmax was not found to be reported in the literature for medrysone and stanozolol as well as the FF concentration for epiandrosterone, stanozolol, dydrogesterone, nandrolone, medrysone, and estropipate.

### Steroids compete for the same binding site as progesterone

To determine whether the strong inhibitory effect of the 10 steroids on P4-induced calcium influx was the consequence of competitive action, we performed cross-desensitization experiments by challenging pre-incubated sperm cells with steroids at 10 µM with increasing doses of P4 (**Fig. 5, A**). A significant reduction in the maximum effect without an important shift in the EC_50_ was observed indicating that steroids reduced the efficacy of progesterone without affecting its potency. In contrast, the same experiment performed with increasing doses of PGE1 revealed a slight shift in the EC50 without a decrease in maximum response indicating that steroids reduce the potency of PGE1 but not its efficacy. (**Fig. 5, B**).

**Figure 5:**
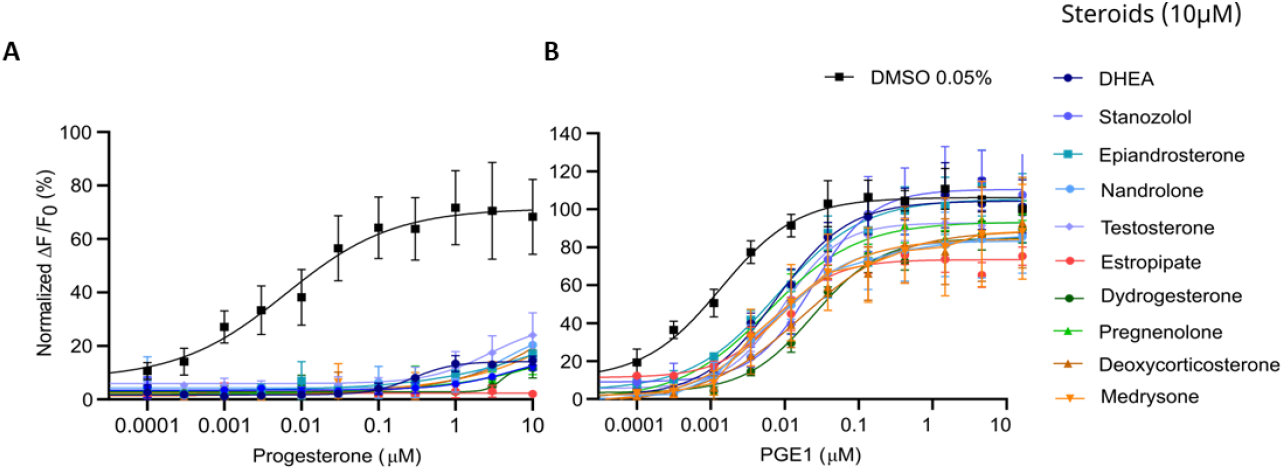
Desensitization experiments with increasing dose of P4 and PGE1 in presence of a fixed dose of steroids. Concentration-response curves of (**A**) the P4-induced and (**B**) the PGE1-induced calcium response in the absence (DMSO) and presence of steroids at 10 µM. Data are presented as the mean of 3 independent experiments with error bars representing the standard error of the mean (SEM).

### Steroids suppress pH-induced activation of CatSper

Besides ligand activation, CatSper can be weakly activated by changes in membrane voltage or alkalization of intracellular pH (pH_i_) (Brown et al., 2019, Wang et al., 2021). Therefore, the 10 selected steroids were evaluated for their ability to affect CatSper-mediated Ca^2+^ influx induced by intracellular alkalization via the weak base NH_4_Cl. All steroids almost completely suppressed the Ca^2+^ signals evoked by NH_4_Cl at 10 mM (**Fig. 6**). We also measured changes in pH_i_ in the presence of the 10 steroids. None of the steroids induced an increase in intracellular pH (**Supplementary Fig. 4**), indicating that CatSper activation by steroids is not triggered by the increase in pH_i_. Our results suggest that the selected steroids inhibit alkaline activation of CatSper, by directly modulating CatSper.

**Figure 6:**
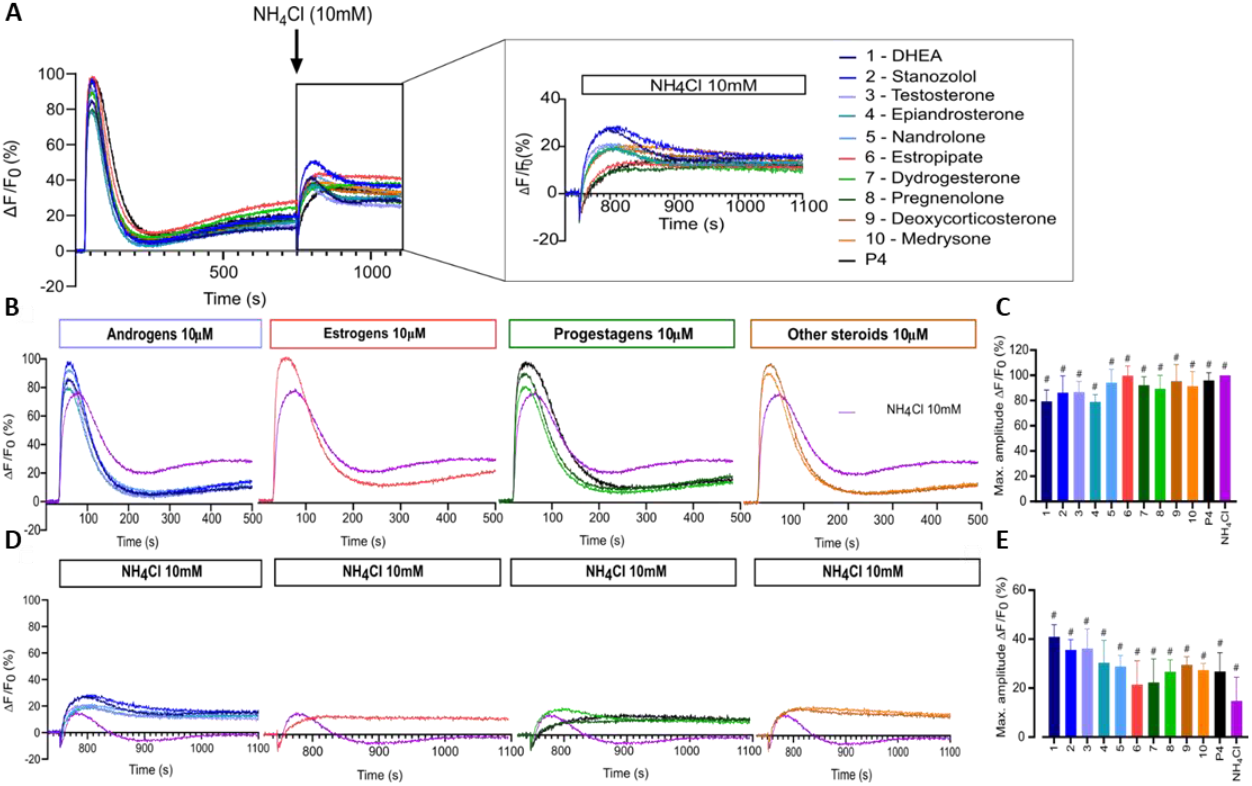
Alkaline-evoked [Ca^2+^]i signals in presence of steroids. (**A**) Overall representative curves of calcium influx after first addition of steroids at 10µM and second addition of NH4Cl at 10mM. (**B**) Representative curves of [Ca^2+^]i induced by steroids at 10µM and positive control NH4Cl at 10mM. (**C**) Bar plots representing maximum amplitude of [Ca^2+^] in presence of steroids and NH4Cl (n=3). (**D**) [Ca^2+^]i increase evoked by single dose of NH4Cl (10mM) in presence of steroids at 10µM. (**E**) Bar plot of maximum amplitude evoked by a single dose of NH4Cl in presence of steroids (n=3), p(#)<0.0001.

### Structure-activity relationship of agonist steroid compounds

In the absence of a resolved structure for human CatSper, we used pharmacophore modelling and machine learning to search *in silico* for additional steroid compounds with a strong activating effect as well as define the steroid structure-activity relationship (SAR) responsible for the CatSper activation. We first constructed a pharmacophore model for the steroids with agonist activity that we identified in our initial screening (**Fig. 2, B & C**), and then combined them with a statistical regression model capable of relating their physicochemical properties to their activity profile. After validating the ability of our combined model to identify compounds that could increase the calcium influx against a blind dataset, we used it to screen 783 compounds *in silico*. Of the 20 top compounds that we selected and tested experimentally, 11 showed activity in the high nanomolar range (**Supplementary Table 1**). Of particular interest are the synthetic steroids S46 and S65 (**Fig. 7, A**). These two compounds potently increased [Ca^2+^]_i_ in human sperm (EC_50_ of 32 nM and 73 nM, respectively) at concentrations close to P4 (EC_50_ of 2.5 nM) (**Fig. 7, B & C**). These two compounds also significantly decreased the response to P4 (IC_50_ of 46 nM and 138 nM, respectively) (**Fig. 7, D & E**). Among the different physicochemical properties we considered for the construction of the statistical regression model to predict the agonist effect of new steroids, we found a significant positive correlation between the predicted IC_50_ value for HERG K^+^ channel block (QPlogHERG) and the number of amide groups and the square of the dipole moment divided by the molecular volume (dip^2^/V). This finding is not unexpected as dip^2^/V is a crucial term in the free energy of solvation of a dipole with volume V. Additionally, dip^2^/V and the octanol/water partition coefficient (QPlogP_o/w_), as well as dip^2^/V and the y-component of the dipole moment, were found to be significantly positively correlated with the dependent variable, in this case, the agonist effect. The coefficients of the properties used in the estimated model can be found in **Supplementary Table 2**. From our model, it is clear that the desolvation of the steroids and their dipole moment in the direction of the electric field inside the channel with respect to their volume are key properties that need to be fine-tuned when designing new steroid analogues that target CatSper. To investigate the SAR of steroid activation of the CatSper channel, we evaluated a wide range of steroids with different functional groups in each of the steroid rings for their ability to induce calcium influx in human sperm. Our results show that the CatSper channel is activated by multiple steroids with various structural modifications. Structural analysis of the steroids provides useful insights into substitutions that can improve the potency of P4 analogues (**Fig. 8**). In particular, various substitutions on the A-ring could be tolerated, including a pyrazole ring (stanozolol, **Supplementary Table 1**), resulting in moderate CatSper activators. Interestingly, the removal of the double bond present on the A-ring of P4 also increased the agonist activity of some compounds such as pregnenolone, S67, S85, and S45 (**Fig. 7, B & C**, and **Supplementary Table 1**). This suggests that some flexibility in this ring may facilitate stronger interactions with the channel residues. In agreement with previous reports (Carlson et al., 2022b), methylation at the C19 position on the B-ring does not appear to be important for the activity, whereas our results indicate that methylation at the C7 position on the same ring can be tolerated. Any significant substitution on the C-ring rendered the steroids inactive or unable to bind at all. Only steroids with a hydroxyl group at the C13 or C14 position seemed tolerable in our dataset. Finally, bulky aliphatic substitutions at the C17 position of the D-ring led to a dramatic decrease in agonist activity, but small substitutions with hydrogen bond donors had a modest activating effect (deoxycorticosterone, stanozolol, norgestimate, **Supplementary Table 1**), suggesting that this part of the steroid may play a key role in stabilizing binding and thus activating CatSper. Notably, structural modifications on the D-ring have been reported to convert agonists to antagonists (Carlson et al., 2022b) and, therefore, this region can be exploited in the design of potent steroidal antagonists of CatSper (**Supplementary Fig. 1**).

**Figure 7:**
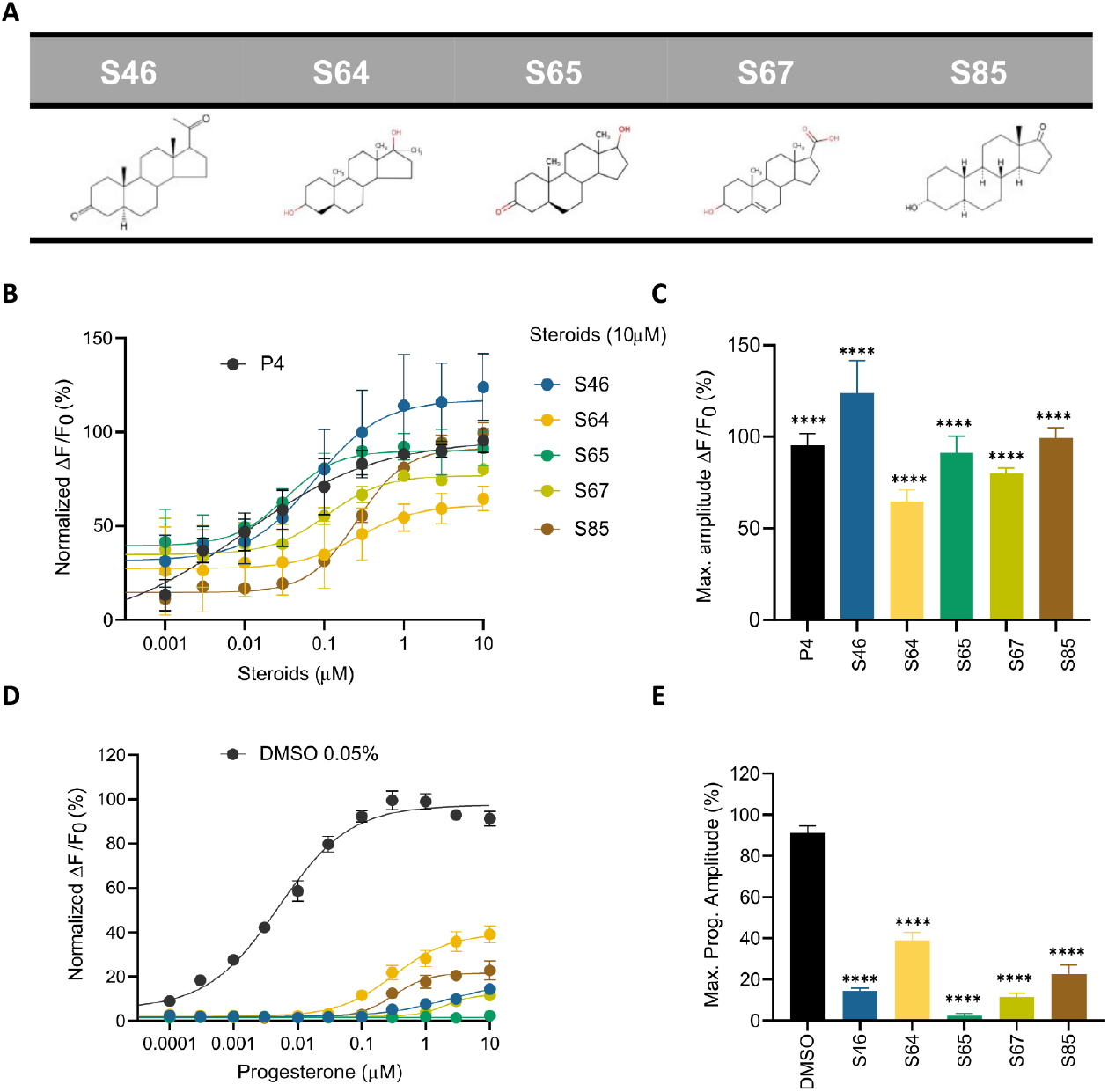
Ca^2+^ response and P4-induced response in presence of steroids predicted by pharmacophore and statistical modeling. (**A**) Chemical structure of the top five predicted steroids selected from the pharmacophore and statistical regression models. (**B**) Concentration-response curves comparing the potencies of the steroids with P4. (**C**) Bar plot of the maximum signal amplitude (ΔF/F0 (%)) of the steroid-induced calcium increase at 10 µM. (**D**) Concentration-response curves comparing the inhibition of P4-induced [Ca^2+^]i influx by the top five predicted steroids at 10 µM, normalized to DMSO. The generated EC50s were the following: EC50 (P4): 0.005 µM, EC50 (S46): 2.254 µM, EC50 (S64): 0.322 µM; EC50 (S65): NA; EC50 (S67): 2 µM; EC50 (S85): 0.32 µM. (**E**) Bar plot of the maximum signal amplitude (ΔF/F0 (%)) of P4-induced [Ca^2+^]i influx in the presence of the top five predicted steroids at 10 µM, normalized to DMSO. All steroids induced a significant inhibition of the response elicited by P4. Data are plotted as the mean of three independent experiments with error bars representing the SD and expressed as a percentage of the response elicited by P4 (10 µM) or DMSO (0.05%); p(****)<0.0001.

**Figure 8:**
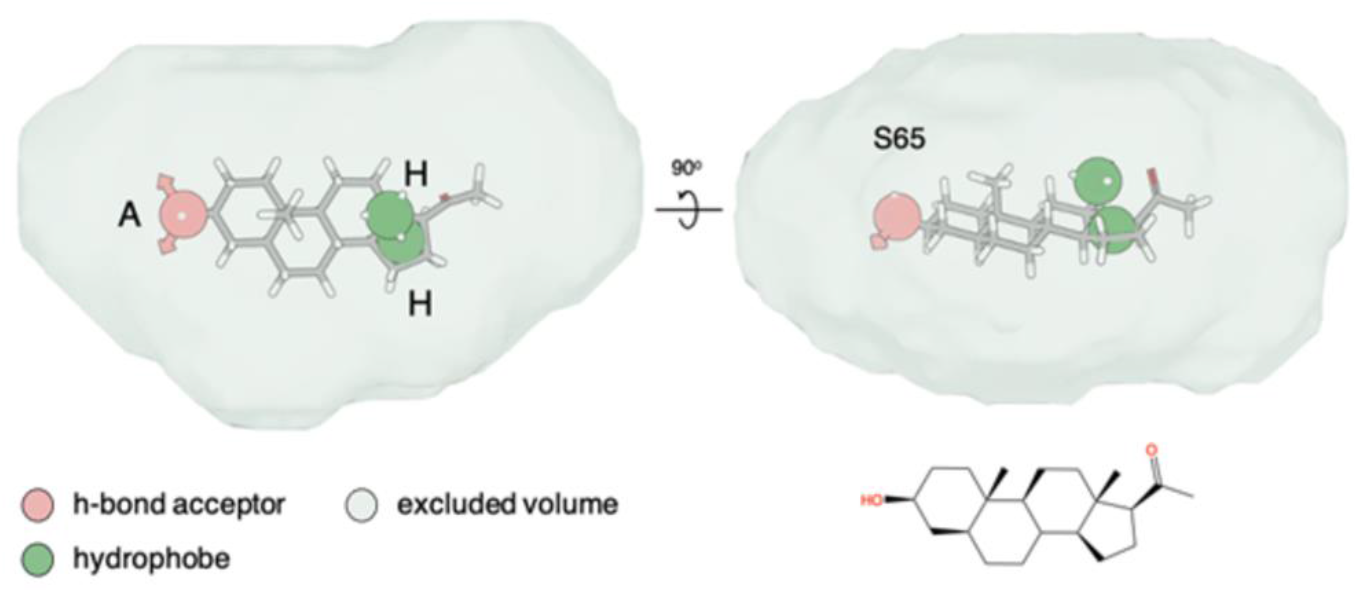
Pharmacophore hypothesis. Top and side view of the fitting of compound S65 into the pharmacophore hypothesis depicting the minimum pharmacophore features that a ligand needs to satisfy for it to interact non-covalently with its receptor. The hydrogen-bond acceptor (“A”) and hydrophobe (“H”) pharmacophore features are shown as spheres, while the excluded volume that was considered based on the shape of the agonist compounds is shown as a surface.

### Stanozolol and estropipate significantly improve sperm penetration in viscous media

We finally investigated the effect of the 10 selected steroids on acrosomal exocytosis in human sperm (**Fig. 9, A**). All steroids showed similar results to negative control DMSO except stanozolol, epiandrosterone, and pregnenolone. These steroids induced acrosomal exocytosis in more than 10% of the cells. We also assessed sperm penetration in a viscous medium using a modified Kremer’s test (**Fig. 9, B**). At a distance of 1 cm, the majority of the steroids tested increased the number of sperms that can swim in viscous media, although only stanozolol and estropipate induced a significant increase in cell number similar to P4. In addition, we evaluated the sperm motility in the presence and absence of the selected steroids at 10 µM using CASA (**Supplementary Fig. 5**). Incubation of spermatozoa with the selected steroids for one hour did not alter total and progressive motility (**Supplementary Fig. 5**). Cell viability was also assessed by flow cytometry of PI incubated cells (**Supplementary Fig. 6**). Our results show that steroids at 10 µM are neither cytotoxic nor affecting cell viability.

**Figure 9:**
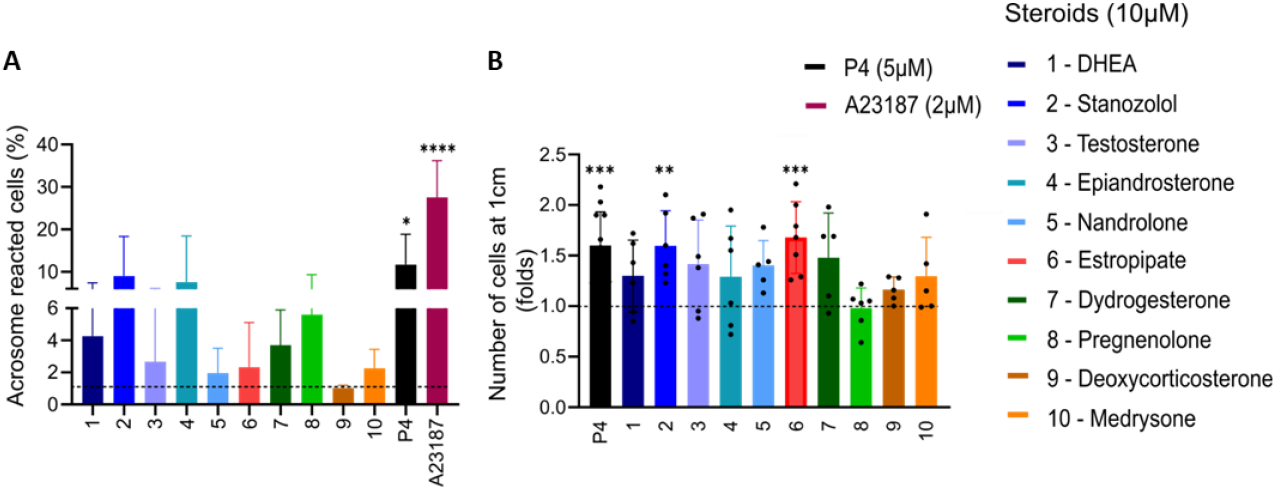
Sperm acrosome reaction and penetration in viscous media induced by steroids. (**A**) Percentage (±SD) of live acrosome-reacted human sperm in the presence of steroids (10 µM), ionophore A23187 (2 µM) used as positive control, normalized to DMSO (shown in dashed black line) (n =5). (**B**) The number of sperm cells that were able to penetrate a glass capillary tubes filled with 1% methylcellulose was assessed after incubation of sperm with steroids at 10 µM for 60 minutes. Number of cells is normalized to DMSO (shown in dashed black line) and results are presented in fold change. Stanozolol and estropipate showed a significant increase in sperm motility in viscous medium (n>5). p(**)<0.01; p(***)<0.001; p(****)<0.0001.

## Discussion

CatSper is the main entry point for Ca^2+^ ions in human spermatozoa, but it is also a promiscuous channel that can be modulated by a wide variety of natural and synthetic molecules (Spehr et al., 2003, Miller et al., 2016, Schiffer et al., 2014, Rehfeld et al., 2016, Tavares et al., 2013, Rahban et al., 2021, Birch et al., 2021, Jeschke et al., 2021). To investigate the impact of diverse chemical structures on [Ca^2+^]_i_, we developed a HTS Ca^2+^ influx assay in human sperm. We screened 1,280 approved and off-patent drugs from the Prestwick chemical library for their potential to either induce a calcium influx or to alter the P4- and PGE1-induced response. We show that a wide range of pharmaceutical compounds belonging to the steroid class, both natural and synthetic, are capable of acting on CatSper. Using a pharmacophore and statistical modelling, we screened *in silico* for steroids that can activate the channel and better understand the molecular characteristics of the steroids that promote CatSper activation.

In this study, we show that the ability of the steroids to activate CatSper is highly variable, with some steroids activating CatSper at low nanomolar concentrations while others are completely ineffective in inducing a calcium response. In particular, 10 steroids had a strong dual activating and inhibiting effect on CatSper and were therefore selected for in-depth analysis of their action. These steroids were either synthetic (stanozolol, estropipate, dydrogesterone, medrysone) or physiological (dehydroepiandrosterone acetate - DHEA -, testosterone, epiandrosterone, nandrolone, pregnenolone, deoxycorticosterone) and were all evaluated for their action on progesterone-, prostaglandin-, pH-activation of human CatSper. In addition to cross-desensitization experiments, our results showed that all 10 steroids directly modulate the CatSper channel and compete for the same binding site as P4, but not PGE1. These observations are consistent with several scientific reports indicating that numerous steroids such as testosterone, hydrocortisone, estradiol, pregnenolone, dihydrotestosterone, and estrone act as true agonists that activate CatSper and increase [Ca^2+^]_i_ in human sperm (Blackmore et al., 1990, Blackmore et al., 1996, Rossato et al., 2005, Luconi et al., 2004, Luconi et al., 1999, Brenker et al., 2018a, Brenker et al., 2018b, Rehfeld, 2020, Jeschke et al., 2021). In particular, a recent study conducted on 15 steroids present in the follicular fluid showed that these steroids cause a rapid increase in calcium influx in human sperm via CatSper (Jeschke et al., 2021). The classical Ca^2+^ response was abolished in CATSPER2^-^/^-^ spermatozoa for all 15 steroids, demonstrating that the response depends on Ca^2+^ influx via CatSper. These results also highlight that steroids and prostaglandins act on distinct binding sites as previously described (Kaupp and Strünker, 2017, Strünker et al., 2011, Lishko et al., 2011, Lishko and Mannowetz, 2018, Miller et al., 2015).

Steroids are widely used as medications for a variety of purposes, some on a long-term basis, and have been available for over 50 years (Bleecker et al., 2020, Kapugi and Cunningham, 2019, Rice et al., 2017). Corticosteroids are one of the most commonly prescribed steroids and are often used to reduce inflammation and suppress the immune system in conditions such as asthma, allergic rhinitis, chronic obstructive pulmonary disease, multiple sclerosis, and others (Bleecker et al., 2020, Ramadan et al., 2019, Rice et al., 2017). Synthetic versions of testosterone and estrogen are also used by some athletes (anabolic steroids) and postmenopausal women, respectively (Cole et al., 2019). Our results suggest that steroids such as deoxycorticosterone significantly increased [Ca^2+^]_i_ and reduced more than 86% of the P4 response with an EC50 = 0.3 µM and an IC50 = 0.5 µM, respectively. Similarly, androgens and estrogens also acted as both potent activators and inhibitors at low nM concentrations. Therefore, an important matter to be addressed here is the physiological significance of the pharmaceutical steroid on CatSper activation. To our knowledge, few previous studies have reported on the concentration of steroids in seminal, oviductal, or follicular fluid. Jeschke et al. reported the average concentration of 18 steroids measured in the follicular fluid of eight ova from four different gonadotropin-stimulated women. Among the 18 steroids, five were steroids that we tested: progesterone, pregnenolone, deoxycorticosterone, DHEA, and testosterone (Jeschke et al., 2021). Here we show that of the 10 steroids analyzed, pregnenolone, dydrogesterone, epiandrosterone, nandrolone, and DHEA are present in sufficient quantities to activate CatSper with EC_02_ below the reported maximum concentration found in blood serum. This suggests that these steroids may interact *in vivo* and potentially affect CatSper activity. It is noteworthy that humans may be exposed to a variety of chemicals that can interfere with CatSper, and these molecules may act additively or synergistically with steroidal compounds to affect CatSper-mediated Ca^2+^ signaling and fertilization in human sperm cells (Brenker et al., 2018a, Rehfeld et al., 2016, Schiffer et al., 2014). To this end, further, *in vitro* and clinical studies are needed to assess the levels of EDCs and other relevant steroids in oviductal fluids and to investigate whether synergistic interactions between these compounds may affect CatSper activity and fertilization. It is worth noting that the antagonizing activity of the steroids on the P4-induced response can also be considered relevant in the field of male contraception (Rehfeld et al., 2022, Wang et al., 2021). Indeed, steroids such as stanozolol and dydrogesterone that strongly reduce the P4 response can serve as a starting point in that direction.

To provide structural insights into the activation of CatSper by steroids, we used the dataset we generated on over 90 steroids and applied pharmacophore modelling and machine learning to define the structure-activity relationship (SAR) associated with the agonist effect of the tested steroids. Due to the wide range of steroids with different functional groups in each of the steroid rings, we were able to refine the SAR for steroid activation of the CatSper channel and provide useful information on substitutions that may enhance the ability of certain steroids to activate CatSper. Our results show that the A-ring of steroids can tolerate various substitutions, including a pyrazole ring, resulting in moderate CatSper activation. Removal of the double bond on the A-ring was shown to increase the activity of some compounds, suggesting that the flexibility of this ring facilitates stronger interactions with the channel residues. Methylation at the C7 position on the B-ring is tolerated, but any significant substitution on the C-ring renders the steroids inactive or unable to bind. Steroids with a hydroxyl group at the C13 or C14 position appeared to be tolerable, and the bulky aliphatic substitutions at the C17 position of the D-ring led to a dramatic decrease in agonist activity. Our results are consistent with those of previous studies that have examined the chemical structures of steroids to determine their affinity for the unknown binding site of CatSper (Carlson et al., 2022a, Carlson et al., 2022b, Jeschke et al., 2021, Xiang et al., 2022). The use of the SAR information allowed us to identify and test several commercially available synthetic steroids in addition to the steroids we initially screened. Some of these steroids were able to increase [Ca^2+^]_i_ and inhibit the P4-induced response at nM concentrations, close to P4. The reduced response suggests that this combined pharmacophore and physicochemical properties modelling approach is effective and allows the identification of steroids capable of activating CatSper already at very low concentrations. The study from Carlson et al. also investigated the effect of multiple steroids and found that CatSper is activated by a wide range of steroids with diverse structural modifications (Carlson et al., 2022b). They examined the modification of residues throughout the steroid skeleton and found that over 30 steroids are capable of activating CatSper. Interestingly, all modifications reduced potency relative to P4, generally without affecting the maximal extent of calcium influx. Here, we have extended this analysis by examining 90 steroidal drugs as well as 30 commercially available synthetic steroids selected by SAR for steroid activation of the CatSper channel. Our data confirm those of Carlson et al. as we found that more than half of the steroids tested can activate CatSper and selectively antagonize P4-induced calcium influx. Furthermore, the potency of activation or inhibition of the P4 response by these steroids is always lower than that of P4. Our *in silico*-based screening identified several steroidal compounds with high potencies capable of inducing Ca^2+^ influx and inhibiting P4-induced response at a low concentration within the nM range (compounds S65 and S46 with EC_50_s of 32 nM and 73 nM and IC_50_s of 46 nM and 138 nM, respectively). In contrast, the most potent steroids inhibiting P4-induced response identified by Carlson et al. such as medroxyprogesterone acetate, levonorgestrel, and aldosterone have IC_50_ values two orders of magnitude higher at 6.6 µM, 32.3 µM, and 33.1 µM, respectively.

In conclusion, our results suggest that diverse steroids act directly on CatSper to activate the channel and compete with P4 for its binding site *in vitro*. Some of the steroids also affect sperm function such as the acrosome reaction and sperm penetration in viscous media. The mechanism of action by which these steroids act on CatSper *in vivo* is complex and remains to be elucidated.

## Data Availability Statement

The original contributions presented in the study are included in the article/supplementary material, further inquiries can be made to the corresponding authors.

## Ethics Statement

Studies involving human participants were reviewed and approved by the Cantonal Ethics Committee of the State of Geneva (CCER #14-147). All donors gave their written informed consent to participate in this study.

## Author contributions

R.R. and S.N. designed the study, L.W. and R.R. coordinated the study. L.W. performed experiments; I.G., L.V. performed *in-silico* modelling; L.W., I.G., L.V., S.G., F.L.G., S.N., and R.R. designed experiments, analyzed and/or interpreted the data, and revised the manuscript critically for important intellectual content, L.W., S.N., and R.R. wrote the manuscript. All authors approved the manuscript.

## Acknowledgments

We thank Gregory Schneiter, Cécile Gameiro, and Jean-Pierre Aubry (F.A.C.S platform, University of Geneva) for technical assistance.

## References

Acosta-Gutierrez, S., Ferrara, L., Pathania, M., Masi, M., Wang, J., Bodrenko, I., Zahn, M., Winterhalter, M., Stavenger, R. A., Pages, J. M., Naismith, J. H., Van Den Berg, B., Page, M. G. P. & Ceccarelli, M. 2018. Getting Drugs into Gram-Negative Bacteriaia: Rational Rules for Permeation through General Porins. ACS Infect Dis, 4, 1487–1498.

Birch, M. R., Dissing, S., Skakkebaek, N. E. & Rehfeld, A. 2021. Finasteride interferes with prostaglandin-induced CatSper signalling in human sperm. Reproduction, 161, 561–572.

Blackmore, P. F., Beebe, S. J., Danforth, D. R. & Alexander, N. 1990. Progesterone and 17α-hydroxyprogesterone. Novel stimulators of calcium influx in human sperm. Journal of Biological Chemistry, 265, 1376–1380.

Blackmore, P. F., Fisher, J. F., Spilman, C. H. & Bleasdale, J. E. 1996. Unusual steroid specificity of the cell surface progesterone receptor on human sperm. Mol Pharmacol, 49, 727–39.

Bleecker, E. R., Menzies-Gow, A. N., Price, D. B., Bourdin, A., Sweet, S., Martin, A. L., Alacqua, M. & Tran, T. N. 2020. Systematic Literature Review of Systemic Corticosteroid Use for Asthma Management. Am J Respir Crit Care Med, 201, 276–293.

Brenker Rehfeld, A., Schiffer, C., Kierzek, M., Kaupp, U. B., Skakkebaek, N. E., StrÜnker, T., SkakkebÆk, N. E. & StrÜnker, T. 2018a. Synergistic activation of CatSper Ca2+ channels in human sperm by oviductal ligands and endocrine disrupting chemicals. Human Reproduction, 33, 1915–1923.

Brenker Schiffer, C., Wagner, I. V., TÜttelmann, F., RÖpke, A., Rennhack, A., Kaupp, U. B. & StrÜnker, T. 2018b. Action of steroids and plant triterpenoids on CatSper Ca2+ channels in human sperm. National Academy of Sciences.

Brenker, C., Goodwin, N., Weyand, I., Kashikar, N. D., Naruse, M., KrÄ Hling, M., MÜller, A., Kaupp, B. & StrÜnker, T. 2012. The CatSper channel: a polymodal chemosensor in human sperm. The EMBO Journal, 31, 1654–1665.

Brown, S. G., Publicover, S. J., Barratt, C. L. R. & Martins Da Silva, S. J. 2019. Human sperm ion channel (dys)function: Implications for fertilization. Human Reproduction Update, 25, 758–776.

Carlson, E. J., Francis, R., Liu, Y., Li, P., Lyon, M., Santi, C. M., Hook, D. J., Hawkinson, J. E. & Georg, G. I. 2022a. Discovery and Characterization of Multiple Classes of Human CatSper Blockers. ChemMedChem, 17, e202000499.

Carlson, E. J., Georg, G. I. & Hawkinson, J. E. 2022b. Steroidal Antagonists of Progesterone- and Prostaglandin E(1)-Induced Activation of the Cation Channel of Sperm. Mol Pharmacol, 101, 56–67.

Cole, T. J., Short, K. L. & Hooper, S. B. 2019. The science of steroids. Semin Fetal Neonatal Med, 24, 170–175.

Jeschke, J. K., Biagioni, C., Schierling, T., Wagner, I. V., Borgel, F., Schepmann, D., Schuring, A., Kulle, A. E., Holterhus, P. M., Von Wolff, M., Wunsch, B., Nordhoff, V., Strunker, T. & Brenker, C. 2021. The Action of Reproductive Fluids and Contained Steroids, Prostaglandins, and Zn(2+) on CatSper Ca(2+) Channels in Human Sperm. Front Cell Dev Biol, 9, 699554.

Kapugi, M. & Cunningham, K. 2019. Corticosteroids. Orthop Nurs, 38, 336–339.

Kaupp, U. B. & StrÜnker, T. 2017. Signaling in Sperm: More Different than Similar. Trends in Cell Biology, 27, 101–109.

Kirichok, Y., Navarro, B. & Clapham, D. E. 2006. Whole-cell patch-clamp measurements of spermatozoa reveal an alkaline-activated Ca2+ channel. Nature, 439, 737–740.

Lishko, P. V., Botchkina, I. L. & Kirichok, Y. 2011. Progesterone activates the principal Ca2+ channel of human sperm. Nature, 471, 387–391.

Lishko, P. V. & Mannowetz, N. 2018. CatSper: a unique calcium channel of the sperm flagellum. Current Opinion in Physiology, 2, 109–113.

Luconi, M., Francavilla, F., Porazzi, I., Macerola, B., Forti, G. & Baldi, E. 2004. Human spermatozoa as a model for studying membrane receptors mediating rapid nongenomic effects of progesterone and estrogens. Steroids, 69, 553–9.

Luconi, M., Muratori, M., Forti, G. & Baldi, E. 1999. Identification and characterization of a novel functional estrogen receptor on human sperm membrane that interferes with progesterone effects. J Clin Endocrinol Metab, 84, 1670–8.

Mannowetz, N., Miller, M. R. & Lishko, P. V. 2017. Regulation of the sperm calcium channel CatSper by endogenous steroids and plant triterpenoids. Proceedings of the National Academy of Sciences of the United States of America, 114, 5743–5748.

Miller, M. R., Mannowetz, N., Iavarone, A. T., Safavi, R., Gracheva, E. O., Smith, J. F., Hill, R. Z., Bautista, D. M., Kirichok, Y. & Lishko, P. V. 2016. Unconventional endocannabinoid signaling governs sperm activation via the sex hormone progesterone. Science (New York, N.Y.), 352, 555–9.

Miller, M. R., Mansell, S. A., Meyers, S. A. & Lishko, P. V. 2015. Flagellar ion channels of sperm: similarities and differences between species. Cell Calcium, 58, 105–113.

Rahban, R., Rehfeld, A., Schiffer, C., Brenker, C., Egeberg Palme, D. L., Wang, T., Lorenz, J., Almstrup, K., Skakkebaek, N. E., Strunker, T. & Nef, S. 2021. The antidepressant Sertraline inhibits CatSper Ca2+ channels in human sperm. Hum Reprod, 36, 2638–2648.

Ramadan, A. A., Gaffin, J. M., Israel, E. & Phipatanakul, W. 2019. Asthma and Corticosteroid Responses in Childhood and Adult Asthma. Clin Chest Med, 40, 163–177.

Rehfeld, A. 2020. Revisiting the action of steroids and triterpenoids on the human sperm Ca2+ channel CatSper. Mol Hum Reprod, 26, 816–824.

Rehfeld, A., Dissing, S. & SkakkebÆk, N. E. 2016. Chemical UV Filters Mimic the Effect of Progesterone on Ca <sup>2+</sup> Signaling in Human Sperm Cells. Endocrinology, 157, 4297–4308.

Rehfeld, A., Frederiksen, H., Rasmussen, R. H., David, A., Chaker, J., Nielsen, B. S., Nielsen, J. E., Juul, A., Skakkebaek, N. E. & Kristensen, D. M. 2022. Human sperm cells can form paracetamol metabolite AM404 that directly interferes with sperm calcium signalling and function through a CatSper-dependent mechanism. Hum Reprod, 37, 922–935.

Ren, D. & Xia, J. 2010. Calcium Signaling Through CatSper Channels in Mammalian Fertilization. Physiology, 25, 165–175.

Rice, J. B., White, A. G., Scarpati, L. M., Wan, G. & Nelson, W. W. 2017. Longterm Systemic Corticosteroid Exposure: A Systematic Literature Review. Clin Ther, 39, 2216–2229.

Rossato, M., Ferigo, M., Galeazzi, C. & Foresta, C. 2005. Estradiol inhibits the effects of extracellular ATP in human sperm by a non genomic mechanism of action. Purinergic Signal, 1, 369–75.

Schiffer, C., MÜller, A., Egeberg, D. L., Alvarez, L., Brenker, C., Rehfeld, A., Frederiksen, H., WÄschle, B., Kaupp, U. B., Balbach, M., Wachten, D., Skakkebaek, N. E., Almstrup, K. & StrÜnker, T. 2014. Direct action of endocrine disrupting chemicals on human sperm. EMBO reports, 15, 758–65.

Spehr, M., Gisselmann, G., Poplawski, A., Riffell, J. A., Wetzel, C. H., Zimmer, R. K. & Hatt, H. 2003. Identification of a testicular odorant receptor mediating human sperm chemotaxis. Science (New York, N.Y.), 299, 2054–8.

StrÜNker, T., Goodwin, N., Brenker, C., Kashikar, N. D., Weyand, I., Seifert, R. & Kaupp, U. B. 2011. The CatSper channel mediates progesteroneinduced Ca 2+ influx in human sperm. Nature, 471, 382–387.

Tavares, R. S., Mansell, S., Barratt, C. L. R., Wilson, S. M., Publicover, S. J. & Ramalho-Santos, J. 2013. p,p’-DDE activates CatSper and compromises human sperm function at environmentally relevant concentrations. Human reproduction (Oxford, England), 28, 3167–3177.

Wang, H., Mcgoldrick, L. L. & Chung, J. J. 2021. Sperm ion channels and transporters in male fertility and infertility. Nat Rev Urol, 18, 46–66.

Xiang, J., Kang, H., Li, H. G., Shi, Y. L., Zhang, Y. L., Ruan, C. L., Liu, L. H., Gao, H. Q., Luo, T., Hu, G. S., Zhu, W. L., Jia, J. M., Chen, J. C. & Fang, J. B. 2022. Competitive CatSper Activators of Progesterone from Rhynchosia volubilis. Planta Med, 88, 881–890.

Zoppino, F. C., Halon, N. D., Bustos, M. A., Pavarotti, M. A. & Mayorga, L. S. 2012. Recording and sorting live human sperm undergoing acrosome reaction. Fertil Steril, 97, 1309–15.

Zou, H. 2006. The Adaptive Lasso and Its Oracle Properties. Journal of the American Statistical Association, 101, 1418–1429.

