## Supplementary data for "The action of physiological and synthetic steroids on the calcium channel CatSper in human sperm"

### Supplementary material

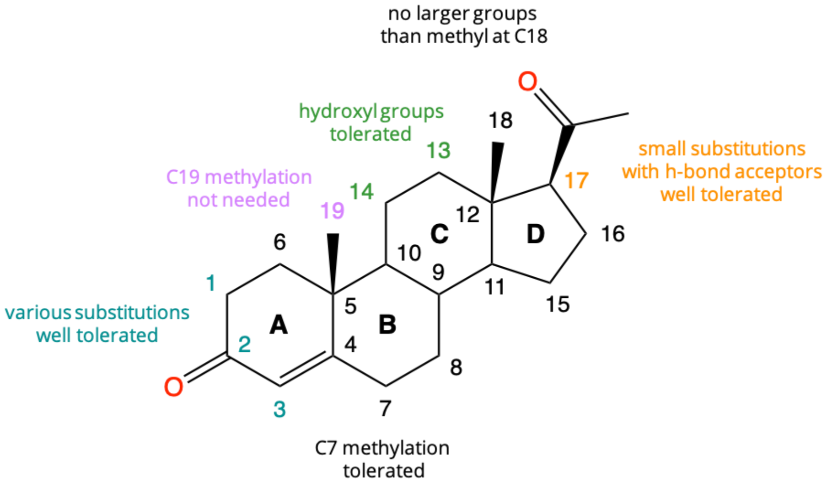


**Supplementary Fig. 1: Structure-activity relationships of CatSper activation by steroids.**

*Pharmacophore and statistical model construction*

The pharmacophore and the statistical modelling was constructed as follow. The original dataset of the 90 compounds was pre-processed using the LigPrep pipeline of the Schrödinger suite (LigPrep, Schrödinger Release 2021, LLC, New York, NY, 2021) at pH=7 and the OPLS4 force field to generate all possible enantiomers and conformers of each steroid in the dataset.

For the construction of the pharmacophore model, compounds that induced more than or equal to 50% increase in [Ca2+]i were labelled as active, while the rest were labelled as inactive. Half of the active compounds (n=12) and all of the inactive compounds (n=70) were first aligned using the steroidal core as the common substructure for alignment. The labelled, pre-aligned set was used to develop several pharmacophore hypotheses using the Phase algorithm of the Schrödinger suite (Dixon et al., 2006). For each hypothesis, an excluded volume shell based only on the active compounds was created to gauge the shape of the binding site. In order to evaluate the different pharmacophore hypotheses generated and to select the most optimal ones, we used the DeepDecoy algorithm (Imrie et al., 2021) to generate a series of decoy compounds (n=932), which match the structure and physicochemical properties of the 20 active compounds we provided. The remaining active compounds (n=8) that were not used for the pharmacophore hypothesis development were mixed with the decoy compounds (decoy/active dataset), and we screened this dataset with the different pharmacophore hypotheses we developed. The top three pharmacophore hypotheses that were able to recover all the active compounds in their top ten hits were selected for further evaluation. To design an appropriate statistical model of the expected agonist effect based on the physicochemical properties, we considered 21 physicochemical properties for each of the 90 compounds of the original dataset. We extended the number of covariates by considering interactions up to the second-degree (i.e. combinations of three variables) between the properties, resulting in a design matrix of 1793 covariates. Based on the distribution of the dependent variable, a regression model with zero-inflated beta distribution (Ospina and Ferrari, 2012) appeared to be suitable. We proceeded to the selection of variables with a penalized regression approach, using the Adaptive-Lasso estimator (Zou, 2006) to select the most relevant covariates. As the relationship between some of the covariates and the dependent variable exhibited clear non-linear relationship patterns, we also included non-parametric smoothing splines for some of the selected covariates in addition to the fixed linear effects. Throughout the iterative development of the model, we prevented overfitting by measuring likelihood-based estimators of prediction error, such as the AIC (Akaike, 1974), and empirical measures of predictive performance, such as the root mean square error (RMSE) estimated by a repeated 10-fold cross-validation. The final model was selected to minimize these criteria. Statistical analysis was performed in the R language using the GAMLSS library for the model estimation (Stasinopoulos, 2017).

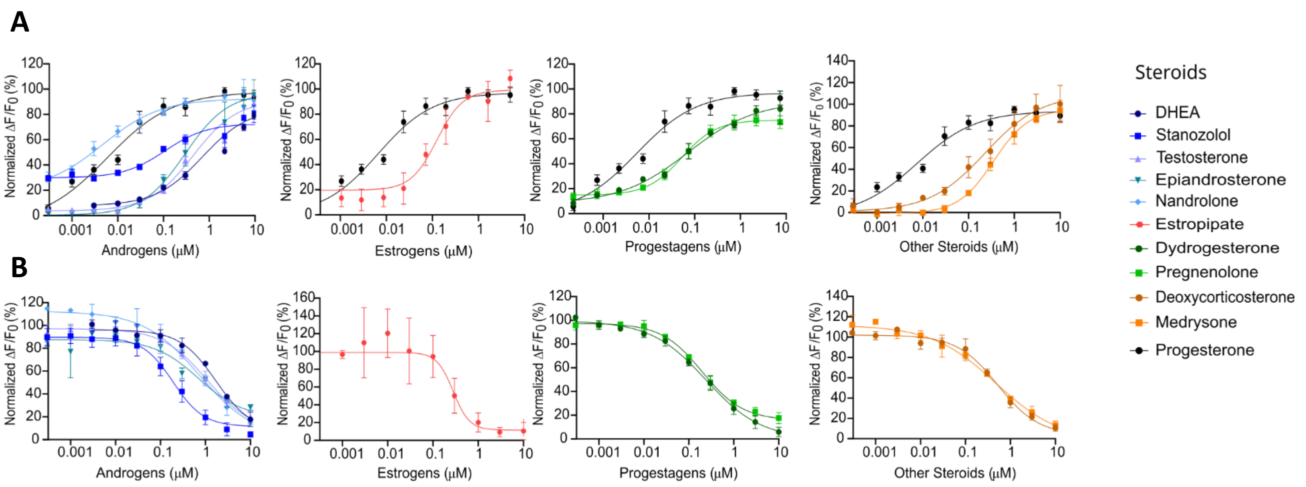


**Supplementary Fig. 2: The 10 selected steroids are capable to induce $Ca^{2+}$ influx and inhibit P4-induced $[Ca^{2+}]_i$ increases in human sperm cells**. (**A)** Cconcentration-response curves of all 10 steroids grouped by steroid classes and compared with the positive control P4 (black curve). (**B)** Concentration-response curves comparing the inhibition of P4-induced $Ca^{2+}$ influx by the 10 steroids grouped by steroid classes. Data are plotted as the mean of three independent experiments with error bars representing the SEM and are expressed as a percentage of the response elicited by the saturating concentration of P4 (10 µM) or DMSO (0.05%).

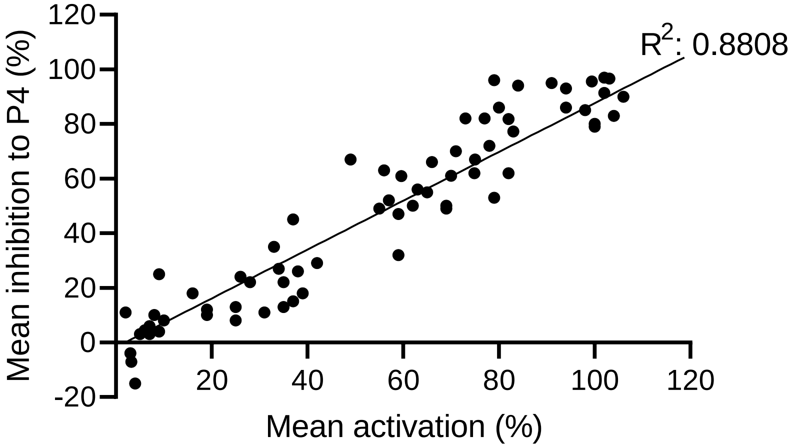


**Supplementary Figure 3: Mean activation of steroids is correlated with P4-inhibition.** Correlation graph of 72 steroids (42 validated steroids from the Prestwick library and 30 additional steroids generated from pharmacophore and statistical regression models). Each dot represents the mean maximal amplitude (%) triggered by individual steroids alone (mean activation) or in presence of P4 (mean inhibition). $R^2$ = 0.8808

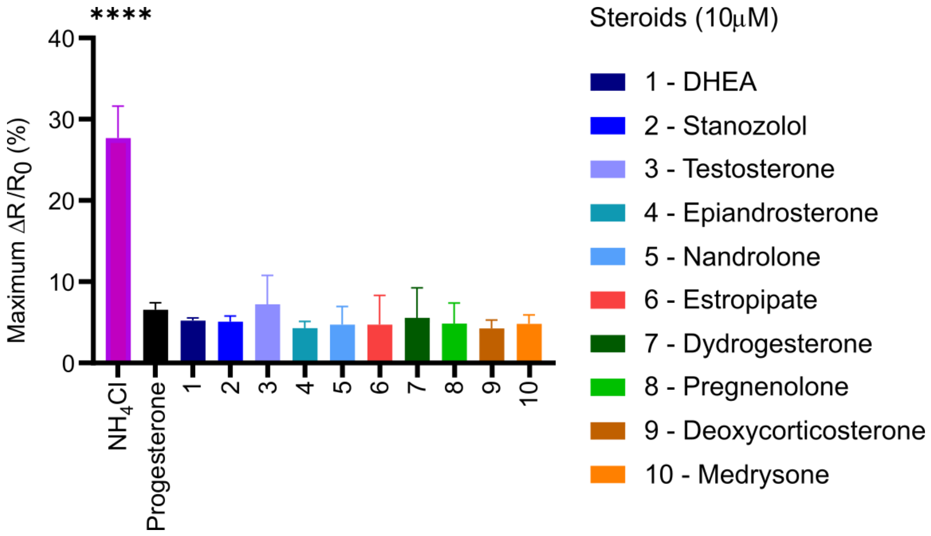


**Supplementary Figure 4: None of the tested steroids induces an increase in intracellular pH.** Changes in intracellular pH ($pH_i$) were measured in sperm loaded with the pH-indicator BCECF-AM (2.5 µM) for 30 minutes at 37°C. Differences in pHi, reflected by changes in the fluorescence ratio, were analyzed as $\Delta R/R_0$ (%) and were monitored before and after the injection of steroids (10 µM). DMSO 0.05% was used as a negative control and $NH_4Cl$ (30mM) as a positive control. Data are presented as the mean of three independent experiments with error bars representing the SEM; p (****)<0.0001.

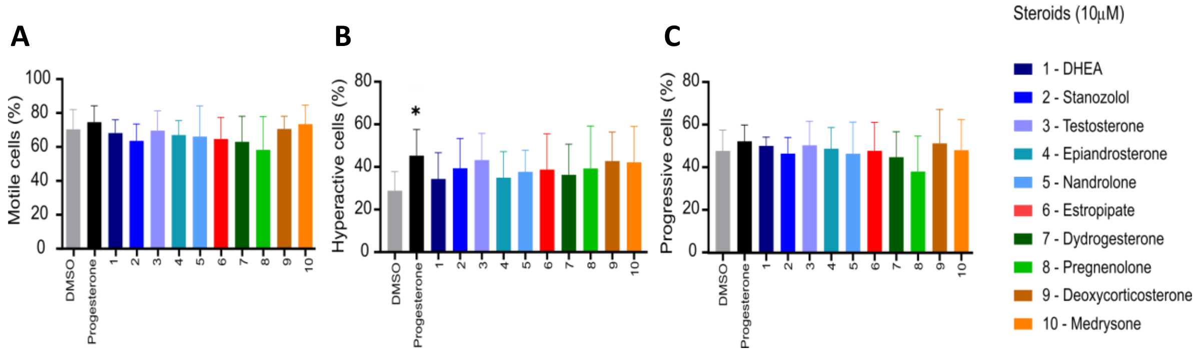


**Supplementary Figure 5: Effect of steroids on motility.** Cells were incubated for 60 minutes with steroids (10 µM) or buffer (DMSO 0,05%) (n≥4). None of the selected steroids induced a significant change in (**A**) overall motility, (**B**) hyperactivation or (**C**) progressive motility. All data were compared with vehicle; p(*)<0.05.

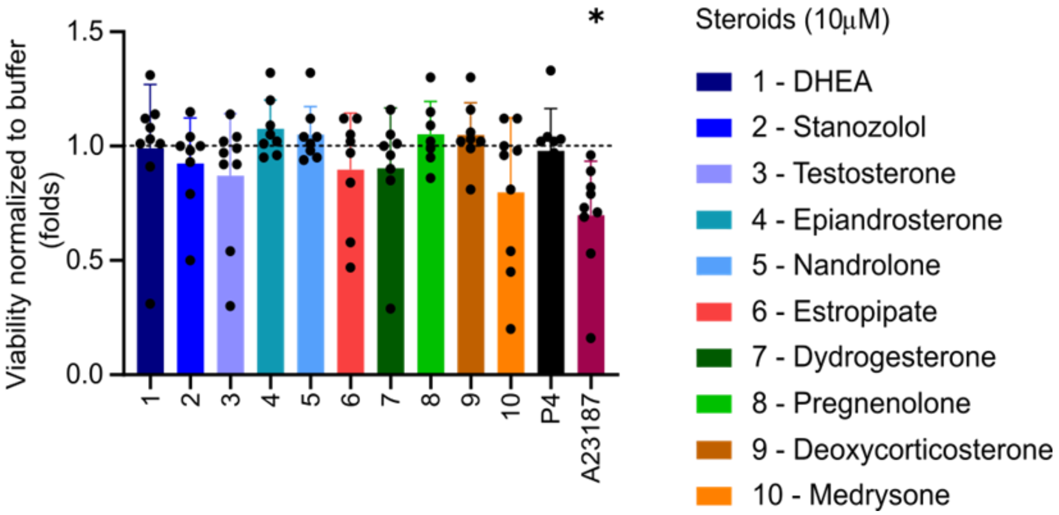


**Supplementary Fig 6: Viability of sperm cells in the presence of the 10 selected steroids.** Cell viability was assessed by flow cytometry of cells incubated with propidium iodide (PI) incubated cells. Cells were incubated in steroids (10 µM) for 60 minutes or positive control ionophore A23187 (2 µm) for 45 minutes. Human sperm viability is not altered in the presence of the steroids at 10 µM. Data are presented as the mean of nine independent experiments with error bars representing the SEM; p(*)<0.05.

| Chemical name | Max amplitude (%) (± SD) | Max P4 inhibition (%) (± SD) | Physiological presence | Mean $EC_{50}$ µM (±SD) | Mean $IC_{50}$ (P4) µM (±SD) | Mean $IC_{50}$ (PGE1) -µM (±SD) |
|---|---|---|---|---|---|---|
| progesterone | **100 (±0)** | **79 (±7)** | Yes | **0.002 (±0.001)** | **0.006 (±0.002)** | **NA** |
| S65 | **91 (±9)** | **95 (±1)** | No | **0.032 (±0.009)** | **0.046 (±0.008)** | **ND** |
| pregnenolone | **73 (±5)** | **82 (±5)** | Yes | **0.070 (±0.016)** | **0.199 (±0.008)** | **0.011 (±0.005)** |
| S46 | **141 (±32)** | **90 (±6)** | No | **0.073 (±0.02)** | **0.138 (±0.03)** | **NA** |
| stanozolol | **79 (±7)** | **96 (±2)** | No | **0.081 (±0.023)** | **0.190 (±0.045)** | **0.117 (±0.046)** |
| dydrogesterone | **84 (±12)** | **94 (±4)** | No | **0.104 (±0.103)** | **0.240 (±0.042)** | **44.84 (±25.54)** |
| S67 | **80 (±3)** | **86 (±3)** | No | **0.108 (±0.03)** | **0.253 (±0.067)** | **ND** |
| estropipate | **106 (±7)** | **90 (±10)** | No | **0.159 (±0.373)** | **0.280 (±0.01)** | **0.282 (±0.373)** |
| epiandrosterone | **78 (±5)** | **72 (±2)** | Yes | **0.207 (±0.033)** | **0.625 (±0.102)** | **NA** |
| medrysone | **104 (±17)** | **83 (±4)** | No | **0.230 (±0.103)** | **0.471 (±0.275)** | **NA** |
| S85 | **99 (±6)** | **96 (±2)** | No | **0.259 (±0.042)** | **0.495 (±0.051)** | **ND** |
| S64 | **65 (±6)** | **55 (±1)** | No | **0.265 (±0.079)** | **0.682 (±0.068)** | **ND** |
| estrone | **49 (±7)** | **67 (±2)** | Yes | **0.267 (±0.016)** | **0.560 (±0.051)** | **NA** |
| S76 | **57 (±6)** | **52 (±3)** | No | **0.277 (±0.006)** | **1.441 (±0.835)** | **ND** |
| S61 | **102 (±6)** | **97 (±1)** | No | **0.28 (±0.68)** | **0.25 (±0.048)** | **ND** |
| deoxycorticosterone | **94 (±12)** | **86 (±3)** | Yes | **0.295 (±0.062)** | **0.523 (±0.081)** | **NA** |
| nandrolone | **98 (±9)** | **85 (±1)** | No | **0.306 (±0.033)** | **0.729 (±0.109)** | **NA** |
| S62 | **82 (±7)** | **82 (±6)** | Yes | **0.33 (±0.078)** | **0.802 (±0.142)** | **ND** |
| formestane | **71 (±4)** | **70 (±8)** | No | **0.353 (±0.088)** | **1.216 (±0.442)** | **NA** |
| S75 | **37 (±3)** | **45 (±6)** | No | **0.358 (±0.182)** | **0.954 (±0.31)** | **ND** |
| S84 | **103 (±8)** | **97 (±2)** | No | **0.39 (±0.108)** | **0.198 (±0.049)** | **ND** |
| testosterone | **87 (±6)** | **81 (±6)** | Yes | **0.407 (±0.062)** | **1.092 (±0.157)** | **NA** |
| S45 | **125 (±18)** | **86 (±10)** | No | **0.445 (±0.098)** | **0.568 (±0.14)** | **ND** |
| exemestane | **63 (±6)** | **56 (±2)** | No | **0.557 (±0.12)** | **0.309 (±1.113)** | **NA** |
| dehydroisoandosterone 3-acetate | **77 (±3)** | **82 (±6)** | Yes | **0.584 (±0.043)** | **1.692 (±0.488)** | **3.704 (±0.961)** |
| fluorometholone | **42 (±1)** | **29 (±12)** | No | **0.702 (±0.329)** | **6.505 (±11.59)** | **NA** |
| estradiol-17 beta | **75 (±8)** | **67 (±3)** | Yes | **0.761 (±0.235)** | **1.451 (±0.225)** | **NA** |
| finasteride | **39 (±5)** | **18 (±6)** | No | **0.780 (±0.348)** | **NA** | **>100** |
| lithocholic acid | **59 (±4)** | **32 (±10)** | Yes | **0.909 (±0.079)** | **0.434 (±0.084)** | **NA** |
| 3-alpha-hydroxy-5-beta-androstan-17-one | **66 (±2)** | **66 (±2)** | No | **0.915 (±0.312)** | **1.517 (±0.216)** | **NA** |
| norgestimate | **79 (±3)** | **53 (±5)** | No | **0.918 (±1.111)** | **8.048 (±1.306)** | **NA** |
| equilin | **62 (±6)** | **50 (±11)** | No | **0.999 (±0.097)** | **1.85 (±0.416)** | **NA** |
| androsterone | **70 (±2)** | **61 (±6)** | Yes | **1.025 (±0.061)** | **2.108 (±0.759)** | **5.383 (±24.22)** |
| oxymetholone | **82 (±12)** | **62 (±9)** | No | **1.299 (±0.423)** | **4.738 (±1.64)** | **11.75 (±3.373)** |
| ethinylestradiol | **69 (±12)** | **50 (±14)** | No | **1.799 (±1.018)** | **3.363 (±1.02)** | **6.678 (±3.62)** |
| S82 | **60 (±10)** | **61 (±3)** | No | **1.906 (±0.026)** | **5.216 (±0.671)** | **ND** |
| S77 | **100 (±6)** | **80 (±2)** | No | **1.969 (±0.393)** | **3.073 (±0.149)** | **ND** |

| | | | | | | |
|---|---|---|---|---|---|---|
| S44 | **102 (±18)** | **91 (±6)** | No | **2.132 (±1.311)** | **1.56 (±0.403)** | **ND** |
| S43 | **83 (±5)** | **77 (±3)** | Yes | **2.213 (±0.966)** | **3.122 (±0.884)** | **ND** |
| fulvestrant | **28 (±4)** | **22 (±6)** | No | **2.242 (±2.254)** | **6.953 (±2.474)** | **NA** |
| corticosterone | **55 (±0)** | **49 (±5)** | Yes | **2.936 (±0.486)** | **6.096 (±1.884)** | **NA** |
| S81 | **33 (±3)** | **35 (±5)** | No | **5.432 (±0.611)** | **12.9 (±15.5)** | **ND** |
| S78 | **38 (±19)** | **26 (±14)** | No | **58.07 (±208.7)** | **8.567 (±6.16)** | **ND** |
| norethindrone | **28 (±12)** | **22 (±5)** | No | **6.053 (±3.101)** | **4.346 (±3.028)** | **NA** |
| S71 | **56 (±10)** | **63 (±6)** | No | **7.023 (±9.242)** | **1.291 (±0.166)** | **ND** |
| canrenone | **69 (±22)** | **49 (±1)** | No | **7.506 (±6.101)** | **NA** | **NA** |
| urosiol | **25 (±8)** | **13 (±3)** | Yes | **7.888 (±2.169)** | **NA** | **12.17 (±6.159)** |
| mifepristone | **9 (±4)** | **25 (±25)** | No | **13.46 (±4.426)** | **10.95 (±5.198)** | **NA** |
| adrenosterone | **26 (±2)** | **24 (±3)** | No | **14.46 (±13.58)** | **30.75 (±342.6)** | **NA** |
| S68 | **37 (±14)** | **15 (±2)** | No | **15.34 (±18.21)** | **NA** | **ND** |
| lynestrenol | **59 (±24)** | **47 (±27)** | No | **17.26 (±5.523)** | **9.360 (±3.302)** | **11.71 (±3.784)** |
| ethynodiol diacetate | **35 (±14)** | **22 (±14)** | No | **17.68 (±5.114)** | **13.95 (±7.309)** | **12.19 (±9.644)** |
| S79 | **35 (±19)** | **13 (±4)** | No | **22.26 (±9.914)** | **NA** | **ND** |
| nomegestrol acetate | **19 (±12)** | **12 (±17)** | No | **22.65 (±7.749)** | **41.17 (±3.135)** | **51.99 (±24.92)** |
| canrenoic acid potassium salt | **34 (±2)** | **27 (±10)** | No | **23.35 (±28.88)** | **23.19 (±3.01)** | **16.34 (±9.27)** |
| chlormadinone acetate | **4 (±2)** | **0 (±15)** | No | **34.71 (±8.298)** | **NA** | **38.44 (±23.34)** |
| S73 | **16 (±2)** | **18 (±1)** | No | **76.54 (±80.74)** | **NA** | **ND** |
| budesonide | **7 (±2)** | **3 (±8)** | No | **NA** | **NA** | **>100** |
| ethisterone | **8 (±5)** | **10 (±9)** | No | **NA** | **5.128 (±60.47)** | **NA** |
| ethynylestradiol 3-methyl ether (mestranol) | **9 (±2)** | **4 (±3)** | No | **NA** | **NA** | **NA** |
| gestrinone | **2 (±2)** | **11 (±21)** | No | **NA** | **33.57 (±18.96)** | **NA** |
| megestrol acetate | **3 (±3)** | **-4 (±11)** | No | **NA** | **NA** | **NA** |
| norgestrel-(-)-d | **5 (±3)** | **3 (±5)** | No | **NA** | **75.56 (±2.899)** | **NA** |
| S80 | **10 (±1)** | **8 (±2)** | No | **NA** | **NA** | **ND** |
| S72 | **94 (±6)** | **93 (±3)** | No | **NA** | **NA** | **ND** |
| S66 | **31 (±18)** | **11 (±10)** | No | **NA** | **NA** | **ND** |
| S70 | **25 (±25)** | **8 (±7)** | No | **NA** | **NA** | **ND** |
| S69 | **19 (±17)** | **10 (±4)** | No | **NA** | **NA** | **ND** |
| S74 | **7 (±2)** | **6 (±8)** | No | **NA** | **NA** | **ND** |
| S83 | **3 (±1)** | **0 (±9)** | No | **NA** | **NA** | **ND** |
| S63 | **6 (±3)** | **4 (±3)** | No | **NA** | **NA** | **ND** |

**Supplementary Table 1: Summary list of the 42 validated steroids and the 30 additional predicted steroids by SAR in order of increasing potency.** Summary table of all 42 validated steroids out of the 90 steroids from Prestwick library, as well as the additional 30 predicted steroids. $EC_{50}$s and $IC_{50}$s are in µM; pooled results from three independent experiments.

| | Estimate | SD | t-value | p-value | |
|---|---|---|---|---|---|
| (Intercept) | 2.0206 | 0.5167 | 3.9104 | 0.0001 | *** |
| QPlogPo/w | 0.2000 | 0.1277 | 1.5660 | 0.1187 | |
| QPlogHERG | 1.1337 | 0.2076 | 5.4598 | 1.2E-07 | *** |
| number of amide groups \| dip^2/V | 42.6134 | 16.7341 | 2.5465 | 0.0115 | * |
| QplogPo/w \| dip^2/V | 6.5199 | 2.7610 | 2.3614 | 0.0190 | * |
| y-component of dipole moment \| dip^2/V | 3.4160 | 1.6682 | 2.0477 | 0.0417 | * |
| number of rotatable bonds \| number of h-bond donors \| dip^2/V | 1.4152 | 0.7905 | 1.7902 | 0.0747 | , |
| number of h-bond donors \| QPlogPo/w \| dip^2/V | -3.5029 | 2.0682 | -1.6937 | 0.0916 | , |
| number of h-bond donors \| x-component of dipole moment \| dip^2/V | 1.3395 | 0.5290 | 2.5325 | 0.0120 | * |
| QPlogPo/w \| z-component of dipole moment \| dip^2/V | 0.8904 | 0.3476 | 2.5615 | 0.0110 | * |
| y-component of dipole moment \| z-component of dipole moment \| dip^2/V | 0.9267 | 0.4408 | 2.1024 | 0.0366 | * |

**Supplementary Table 2. Estimated parameters and standard errors (SD) of fixed effects parameters of the beta zero inflated semi-parametric model.** Variables whose interaction has been considered in the model are denoted with a vertical line. p(***)<0.001, p(*)<0.05, p(.)<0.1, p(,)<1.
